# Multifunctional CNT-based Platform for Enhanced Dopaminergic Differentiation and Electrochemical Dopamine Sensing from Human iPSC-derived Neurons

**DOI:** 10.64898/2026.11.20.768200

**Authors:** Nerea Pascual Frías, Giuseppe Misia, Lara Rodríguez Sánchez, Claudio Evangelisti, Maurizio Prato, Nuria Alegret, Francisco Gil Bea, Alessandro Silvestri

## Abstract

Pairing neuro-electrochemical sensors with physiologically relevant neuronal models represents a new frontier in the study of neurodegenerative diseases. In this work, we reported an electroactive platform for *in situ* detection of dopamine, combining human iPSC-derived neurons with carbon nanotubes. In this system, carbon nanotubes served a triple role: enhancing the electroactive area, catalyzing dopamine oxidation, and promoting subtype-specific dopaminergic differentiation. We developed a robust protocol achieving functional dopaminergic neurons, confirmed via qPCR, immunocytochemistry, MEA electrophysiology, and calcium imaging. CNT substrates were associated with a significant increase in the TH/neurofilament overlap (37.90 ± 3.80% vs. 26.01 ± 3.75% in controls). Applying differential pulse voltammetry and chronoamperometry, we quantified extracellular dopamine at 1.25 µM and confirmed the stimulus-dependent release induced by benztropine and KCl depolarization. Post-electrochemical immunostaining confirmed that neurons retained their morphology and TH expression. Overall, this work establishes a novel framework for neurodegenerative disease modeling that recapitulates key electrophysiological and neurochemical features of dopaminergic neurons, representing a powerful tool for disease modeling, drug screening, and the development of next-generation neuro-biosensors.

## 1. Introduction

The development of neurodegenerative diseases is frequently associated with a dysregulation of neurotransmitter levels within the central nervous system.^[1–3]^ Dopaminergic (DAergic) neurotransmission, in particular, and associated neural circuits are implicated in a wide range of neurological and neuropsychiatric disorders, including addiction, depression, schizophrenia, and Parkinson’s disease (PD).^[4,5]^ PD is characterized by motor impairments, such as resting tremors and bradykinesia, which arise from the progressive degeneration of DAergic neurons in the *substantia nigra*.^[6,7]^ Patients with PD typically exhibit a loss of 60-80% of their DAergic neurons^[8]^ concurrently with the loss of norepinephrine (NEP)-producing neurons in the *locus coeruleus*.^[9]^ The dopamine (DA) deficiency in the *striatum* disrupts the intricate balance between the direct and indirect pathways of the basal ganglia, which is fundamental to motor control.^[10]^

Current pharmacotherapies, such as levodopa combined with carbidopa and entacapone,^[11]^ or amantadine, which promotes DA release and inhibits its reuptake,^[12]^ provide symptomatic relief but lose efficacy as the disease progresses, highlighting the need for new therapeutic strategies. Accelerating drug discovery for neurodegenerative diseases necessitates the development of high-throughput *in vitro* screening platforms that enable real-time, non-invasive characterization of complex neuronal systems. Current drug screening protocols primarily rely on biochemical assays,^[13]^ high-performance liquid chromatography (HPLC),^[14]^ or microelectrode arrays (MEA).^[15]^ However, the use of HPLC and enzyme-linked immunosorbent assay (ELISA) is limited by high cost, labor-intensive procedures, and the fact that they are end-point analyses, which preclude real-time monitoring of DA release from living neurons. MEA recordings, in turn, report neuronal electrical activity but cannot distinguish between specific neurotransmitters or neuronal subtypes.

Unlike electrophysiological recordings, electrochemical recordings offer a high degree of selectivity between different neuron populations and allow for the real-time, non-destructive detection of neurotransmitters. This capability has led to the development of “brain-on-a-chip” devices, which have garnered significant attention not only for drug screening but also for monitoring stem cell differentiation, evaluating differentiation protocols, and developing new stem cell-based therapies.^[16,17]^ Despite their promise, several key challenges must be addressed for electrochemical recordings to be widely adopted in complex neuronal cultures. The neurotransmitters of interest are typically present at low concentrations, which requires platforms with low limits of detection. Additionally, culture media frequently contain compounds with concentrations up to 350,000 times higher than the target neurotransmitter.^[17]^ This, along with the high probability of electrode fouling, significantly compromises the selectivity and sensitivity of the method. Selecting the most representative cell culture model to mimic the brain’s complex cellular environment accurately remains another crucial challenge. For example, the PC12 Adh cell line is a widely used DAergic model for “brain-on-a-chip” studies.^[16–22]^ However, it does not fully resemble the physiological properties of human primary neurons. The materials used for the electrodes are also critical, as cells are cultured directly on the device. These materials must be biocompatible while supporting the adhesion, connectivity, and maturation of the neurons.^[23,24]^

Carbon nanotubes (CNTs) offer significant potential in neural cell research.^[25,26]^ Their morphology, resembling small neuronal extensions, promotes neuronal growth and neurite outgrowth,^[27,28]^ they are biocompatible,^[29,30]^ and their electrical conductivity provides an effective interface with electroactive cells, increasing synaptic activity and network excitability ^[28,31]^.

However, their role in subtype-specific neuronal differentiation remains largely unexplored. Most studies focus on general neuronal maturation or employ immortalized cell lines such as PC12 or SH-SY5Y, where DAergic features are already partially established.^[32–34]^ As a result, whether CNT-based substrates can influence differentiation toward specific neurotransmitter subtypes in human stem cell-derived systems remains an open question. Furthermore, CNTs are highly effective for the electrochemical detection of DA because they significantly improve the charge and mass transfer kinetics of the electrochemical reaction.^[35,36]^ This property results in superior voltammetric peaks characterized by higher signal intensity and a narrower full width at half maximum (FWHM), directly translating to enhanced sensitivity and selectivity. Additionally, the surface of CNTs can be readily functionalized to tune their physicochemical properties.^[37]^ In both *in vivo* and *in vitro* applications, electrodes modified with CNTs have demonstrated a remarkable increase in DA sensitivity.^[38,39]^

This work introduces a novel electroactive cell cultivation platform capable of electrochemically quantifying DA from living DAergic neurons without altering culture conditions. The platform is based on CNTs, which perform a triple function: (i) to act as a conductive nanomaterial, increasing the sensor’s electroactive area; (ii) to serve as an electrocatalyst, facilitating the electrochemical oxidation of DA; and (iii) to foster neuronal growth and interconnection. By combining CNTs with chemically induced neuronal differentiation, we generated cultures enriched in DAergic and glutamatergic (GLUTergic) neurons, respectively. We used these two neuronal cultures as positive and negative controls to validate the platform. The system successfully detected DA in a linear range from 0.8 to 7 µM, which encompasses the estimated 1.25 µM concentration secreted by the developed DAergic model. We also proved that the platform could distinguish the effect of drugs like benztropine on extracellular DA concentration, confirming its feasibility for drug screening. While previous CNT-based sensing platforms have primarily relied on immortalized cell lines, here we establish a platform compatible with human iPSC-derived neurons, enabling a more relevant biological context for neurochemical sensing. Beyond sensing, we show that CNT-based substrates promote subtype-specific neuronal differentiation, suggesting a previously unexplored role of these materials in influencing the neuronal subtype composition of human iPSC-derived cultures. This integrated approach enables a direct coupling between neuronal development and *in situ* electrochemical readout, providing a new framework for studying neurotransmitter dynamics in human-derived neuronal systems.

## 2. Results and Discussion

### 2.1. Design and electroanalytical performance of the electroactive cell culturing platform

The electroactive cell culturing platform was designed to perform electrochemical and confocal fluorescence microscopy on the same neuronal cell culture, enabling the direct correlation of electrochemical and optical data. As shown in **Figure 1A**, the platform was fabricated using a 0.75 cm x 1.5 cm indium tin oxide (ITO)-covered glass coverslip. The specific thickness of the coverslip (0.16 mm) was chosen to facilitate electrochemical and confocal microscopy measurements on the same sample. A 0.75 cm x 0.75 cm 3D-printed polylactic acid (PLA) cubic well was then attached to the electrode surface with polydimethylsiloxane (PDMS) to define the electroactive and cell culture area. This area was subsequently modified with a double layer of CNTs (diameter = 30 ± 8nm). The first layer consisted of pristine CNTs, while the second layer was composed of CNTs decorated with small gold nanoparticles (average Au NPs dimension = 2.57 ±0.54 nm) named Au-CNTs (**Figure S1**), chosen for their electrocatalytic activity for DA oxidation. Au-CNTs were prepared following a previously published protocol from our group.^[36]^

**Figure 1.**
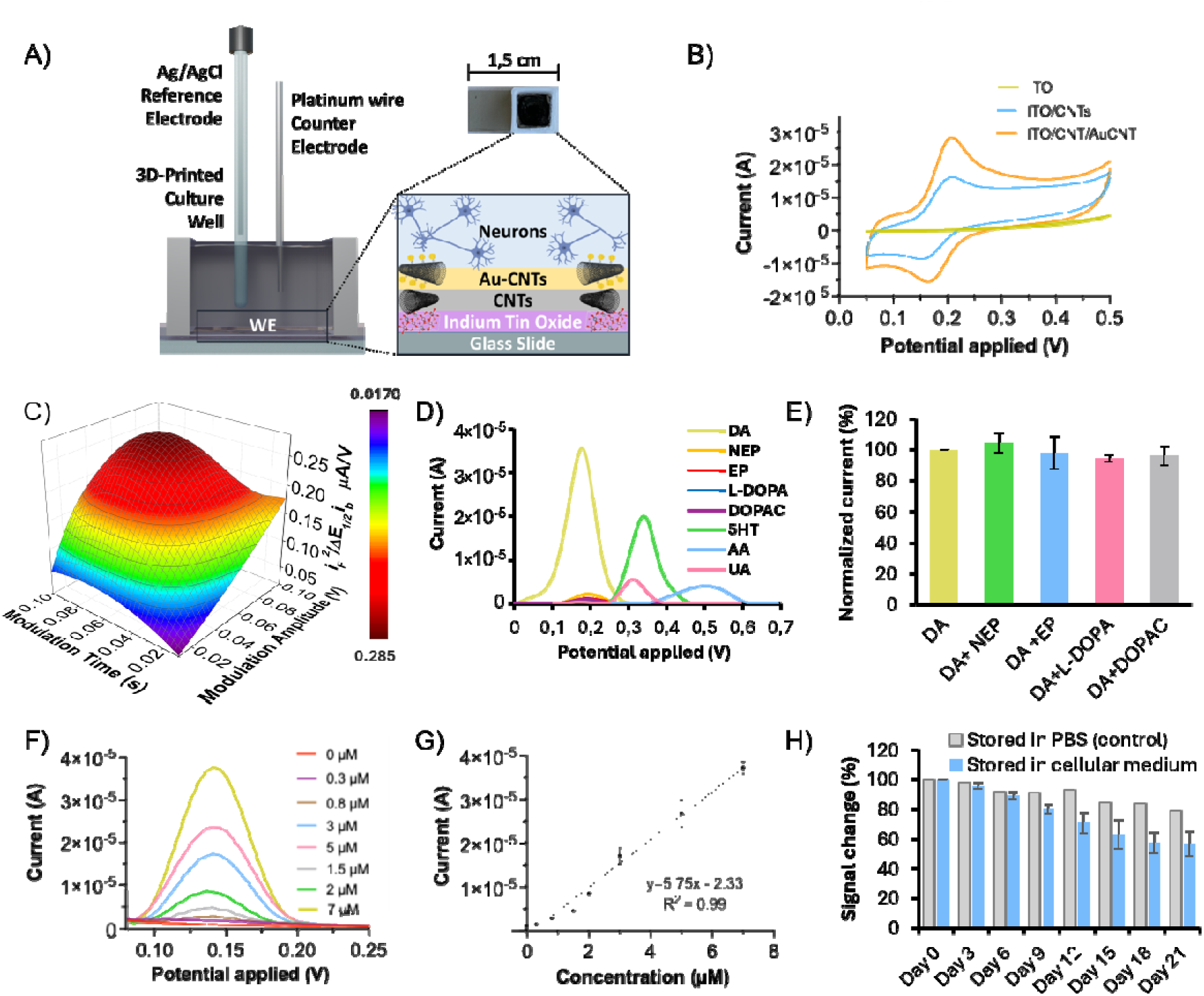
Design and electrochemical performance of the electroactive cell culture platform. A) Schematic representation of the electroactive cell culture platform designed for DA detection at the cellular level. B) CV response to a 100 µM DA solution in Krebs-Ringer buffer, recorded with an ITO electrode at different modification steps: ITO (yellow), ITO/CNT (blue), and ITO/CNT/Au-CNT (orange). C) Response surface of the polynomial function describing the dependence of FoM on MA and MT obtained from the DoE. D) DPV response of an ITO electrode modified with CNT/Au-CNT to 100 µM solutions of DA, 5-HT, UA, AA, EP, NEP, L-DOPA, DOPAC in Krebs-Ringer buffer. E) Interference studies using mixed 1:1 solutions containing 50 µM of DA and 50 µM of each overlapping interferent (EP, NEP, L-DOPA, and DOPAC) in Krebs-Ringer buffer. F) DPV response of an ITO electrode modified with CNT/Au-CNT to different DA concentrations in Krebs-Ringer buffer, namely 0, 0.3, 0.8, 1.5, 2, 3, 5, 7 µM (linear regression was calculated considering DA levels between 0.8 and 7µM). G) Calibration line representing the maximum current of the DPV curves versus DA concentration. H) Normalized DPV current response (signal change, %) toward 100 µM DA (in Krebs-Ringer medium) after continuous incubation at 37 °C in cellular medium (n=3) compared against a control electrode incubated in protein-free PBS solution. Error bars represent the standard deviation. Results are expressed as the average ± SD (n=3).

Briefly, the synthetic protocol proceeds through three main steps. The first involves a single-electron transfer reaction between CNTs and a sulfur-bearing aryl diazonium salt, 4,4′-dithiodibenzenediazonium tetrafluoroborate. The reaction leads to the formation of an aryl radical that binds to the CNTs’ surface. In the second step, the disulfide bond is cleaved, resulting in free thiol moieties on the surface. This addition-cleavage functionalization strategy is crucial because it protects the thiol groups from potential side reactions during the chemical modification while also allowing for control of the oligomerization degree via the reductive cleavage. Finally, the available thiol groups are leveraged to decorate the CNTs with size-controlled Au NPs obtained by metal vapor synthesis (MVS). A significant advantage of these pre-formed nanoparticles is that they consist solely of metal in its reduced form (**Figure S2**) and are weakly stabilized only by the synthesis solvent, without any additional ligands.^[41]^ These properties, coupled with their small size and vast exposed surface area, make the particles ideal catalysts^[42]^ and electrocatalysts.^[43]^ This procedure ultimately allows for stable, homogeneous, and controlled decoration of the CNTs with an Au/C ratio of 11.5 % by ICP-MS and 12.4 % by TGA.^[36]^ For comprehensive characterization of the material, we refer to Misia et al.^[36]^

**Figure S3** reports the morphological and elemental characterization of the successive modification steps of the ITO electrode, namely ITO, ITO/CNT, ITO/CNT/Au-CNT. SEM images acquired after each step illustrate the progressive evolution of the electrode surface. Following the final modification with Au-CNTs, homogeneously distributed bright spots corresponding to gold are clearly visible across the surface. Concurrently, EDX spectra confirm the emergence of a well-defined Au peak, yielding an overall surface Au/C ratio of 1.2% for the final ITO/CNT/Au-CNT electrode.

Cyclic voltammetry (CV) measurements in the presence of 100 µM in Krebs-Ringer buffer (**Figure 1B**) revealed that the CNT/Au–CNT double layer substantially improved oxidation relative to both bare ITO and the CNT-modified electrode. Pristine ITO showed a negligible and highly irreversible oxidation response, indicating that oxidation is strongly disfavored on the bare substrate. Modification with the primary CNT layer led to well-defined redox peaks (E_cat_ = 157 mV, E_an_ = 216 mV, ΔE = 59 mV), reflecting a marked improvement in electrochemical reversibility. The subsequent deposition of the Au–CNT top layer approximately doubled the oxidation peak current and reduced ΔE to 39 mV (E_cat_ = 167 mV, E_an_ = 206 mV), consistent with accelerated electron-transfer kinetics and superior electrocatalytic activity.

Electrochemical impedance spectroscopy (EIS), performed in 100 µM DA in Krebs-Ringer buffer at E_1/2_ = 0.18 V vs. Ag/AgCl (**Figure S4A**, **B**), provided further insight into the origin of these improvements. Pristine ITO exhibited an extremely high charge-transfer resistance (R_CT_), which explains its weak CV response. The primary CNT layer reduced R_CT_ by more than two orders of magnitude, while the Au–CNT layer further lowered it to 1.7 kΩ, in agreement with the narrower peak separation and faster kinetics observed in CV. The electrochemically active surface area (ECSA) was evaluated from [Ru(NH_3_)_6_]^3+^ CVs recorded at multiple scan rates (**Figure S4 C–F**). Whereas the primary CNT layer caused no significant change in ECSA compared with pristine ITO (0.85 cm²), the Au–CNT layer nearly doubled the effective area. This increase aligns with the two-fold enhancement in oxidation current, indicating that the Au–CNT layer contributes to the signal enhancement through both surface-area enlargement and electrocatalytic effect.

Taken together, these results clarify the rationale for the double-layer architecture: the initial CNT layer overcomes the severe charge-transfer resistance of the substrate, while the Au– CNT layer enhances both the available surface area and the electrocatalytic kinetics required for sensitive DA detection. Overall, the use of this double layer induces an increase in faradaic current, improving the sensitivity and signal-to-noise ratio of the platform, while the shift to a lower oxidation potential helps avoid interference from other electroactive molecules like uric acid (UA), ascorbic acid (AA), or other neurotransmitters like serotonin (5HT), which oxidize at higher potentials.^[17]^

We chose differential pulse voltammetry (DPV) as the analytical technique for DA quantification due to its ability to minimize background capacitive current, which is particularly beneficial for modified nanostructured electrodes.^[36]^ This technique significantly improves the sensitivity, signal-to-noise ratio, and limit of detection (LOD) of the final sensor.^[44,45]^

To effectively minimize the capacitive current, we optimized key operational parameters that influence the technique’s performance. Following a previously established Design of Experiments (DoE) from our group,^[36,46]^ we optimized the modulation amplitude (MA), which is the intensity of each potential pulse, and the modulation time (MT), which is the duration of the pulses. By applying a factorial design, we tested all nine possible combinations of three equally spaced levels for each parameter. We ran the experiments in triplicate, using the same electrode for each series and randomizing the sequence to minimize systematic errors.

As described in Equation 1, the DoE aimed to maximize a figure of merit that balances sharp peak resolution, high current intensity, and a high signal-to-noise ratio.

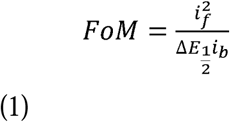

Where *i_f_* is the faradaic current samples at the maximum of the peak, is the baseline current sampled immediately before the start of the oxidation peak and 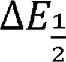 is the width of the peak at medium height.

The average values for each selected level were determined from this final function (**Figure S5**), allowing us to generate the response surface shown in **Figure 1C**. From this response surface, we interpolated the optimal values of 0.052 s for MT and 0.062 V for MA, which were used for all subsequent sensing experiments. For a comprehensive description of the DoE, please refer to the Supporting Information.

Using the optimized DPV parameters, we evaluated the sensor’s selectivity for DA against several electroactive biological interferents, including AA, UA, 5-HT, epinephrine (EP), NEP, levodopa (L-DOPA), and 3,4-dihydroxyphenylacetic acid (DOPAC), all tested at 100 µM. As shown in **Figure 1D**, the voltammetric response for DA was well-resolved from the oxidation peaks of 5-HT, UA, and AA. Instead, catecholamine derivatives sharing a high structural similarity with DA (EP, NEP, DOPAC, and L-DOPA) were oxidized at a similar potential. However, the ITO/CNT/Au-CNT sensor displayed a significantly elevated current response for DA, which was approximately one order of magnitude higher than these structurally related competitors.

To further investigate these overlapping signals, interference studies were performed by analyzing mixed solutions containing 50 µM each of DA and the competing catecholamines via DPV. The resulting interference coefficients, reported in **Figure 1E**, demonstrate that the presence of EP, NEP, L-DOPA, and DOPAC exerts a negligible effect on the DA signal, within experimental error. Overall, these findings confirm that the platform maintains robust selectivity for DA even when challenged by closely related catecholamine competitors.

To elucidate the mechanism underlying this catalytic selectivity toward DA, scan-rate-dependent voltammetry and Tafel analysis were conducted. At a low scan rate (5 mV s^-1^), DA exhibits quasi-reversible behavior with the narrowest peak potential separation (ΔE = 51 mV), indicating rapid electron transfer at the Au-CNT surface. In contrast, EP, NEP, L-DOPA, and DOPAC displayed higher ΔE values characteristic of irreversible behavior (**Table S4**). Increasing the scan rate revealed growing reduction peak intensities for both DA and NEP (**Figure S6**), which is characteristic of an electrochemical-chemical (EC) mechanism where electrochemical oxidation is coupled with rapid intramolecular cyclization to form dopaminochrome and noradrenochrome, respectively. This chemical consumption shifts the equilibrium toward the products, thereby lowering the oxidation overpotential and thermodynamically favoring the oxidation of these two neurotransmitters.

Tafel plots derived from the mass-transport-free regions quantified the charge-transfer kinetics for the different neurotransmitters (**Figure S7**). Notably, DA yielded the lowest Tafel slope (68.15 mV dec^-1^), confirming the fastest electrocatalytic oxidation kinetics on the nanocomposite. The slopes increased for EP and NEP, while L-DOPA and DOPAC exhibited severely sluggish kinetics. Collectively, these findings demonstrate that the catalytic specificity for DA arises from a combination of favorable thermodynamics (a low ΔE and a driving EC mechanism) and rapid charge-transfer kinetics.

Because pulsed voltammetric waveforms like DPV are highly sensitive to electron transfer kinetics, parameters optimized for one neurotransmitter are often suboptimal for another. In this study, the DPV waveform parameters were tuned specifically to match the fast kinetics of DA. Consequently, slower-reacting analogues (such as NEP, EP, DOPAC, and L-DOPA) yield lower signals, thereby enhancing the analytical selectivity of our sensor.

To evaluate the sensor’s performance under optimized conditions, we performed DPV measurements at different DA concentrations in the Krebs-Ringer buffer (**Figure 1F**). We selected this buffer as the electrolyte for our measurements, both in solution and within the neuronal cultures, to prevent interference from the complex compounds typically found in cell culture media. This buffer has a simple composition, containing only inorganic electrolytes (NaCl, KCl, MgCl_2_, CaCl_2_), along with glucose, sucrose, and HEPES. Crucially, it still provides all the necessary nutrients to keep the neurons viable throughout the experiment.

The DPV peak currents were plotted against the DA concentration, demonstrating a linear response in the range of 0.8–7 µM (**Figure 1G**). The platform demonstrated an LOD of 412 nM, with a normalized sensitivity of 7,67 μA cm□² μM□¹. To contextualize these performance metrics, **Table S5** provides a detailed comparison with related Au-CNT composite sensors reported in the literature.

The reproducibility of the measurements was assessed by comparing the linear responses across three distinct electrodes (**Figure S8A**). The results showed a relative standard deviation of 10.3%, confirming that the fabrication protocol provides acceptable batch-to-batch reproducibility. Repeatability was assessed by comparing the linear responses across three separate measurements on the same electrode (**Figure S8B**), showing a relative standard deviation of 4.1%, indicating high consistency.

Although minor signal carryover from adsorbed DA or polyDA fouling was observed after the first repetition of the dose-response measurements (**Figure S**9), this contribution was minimized by buffer washing paired with an electrochemical regeneration protocol (5 CV sweeps between 0 and 0.5 V at 50 mV s^-1^).

While utilizing a Krebs-Ringer buffer minimizes immediate chemical interference, the sensor must undergo prolonged contact with complex media (1:1 DMEM:F12 / Neurobasal-Brainphys) during cell colonization and differentiation, potentially inducing surface biofouling. To evaluate the long-term stability and biofouling behavior of the modified electrode, a 21-day stability study was performed by incubating ITO/CNT/Au-CNT electrodes in the culture medium at 37 °C (**Figure 1H**). Every 3 days, DPV responses to 100 µM DA in Krebs-Ringer buffer were recorded. The signal in cell medium gradually decreased before stabilizing at 56% of its initial value by Day 21.

To distinguish matrix-induced biofouling from baseline electrode degradation, electrodes were incubated in PBS for 21 days at 37 °C. This control accounted for signal losses arising from physical aging of the ITO or nanocomposite, mechanical handling, and polyDA accumulation during repeated measurements. Of the total 44% signal reduction observed in cell culture medium, 21% also occurred in PBS, indicating that only 20% of the loss was attributable to the biological matrix.

To identify the origin of the signal reduction after prolonged incubation, the electrodes were characterized before and after 21 days in cell culture medium. The electrochemical response was assessed using both the outer-sphere redox probe [Ru(NH_3_)_6_]^3+^ and DA (**Figure S10A-C**). In both cases, CVs showed a higher capacitive background, an approximately 20% reduction in faradaic current, increased ohmic drop, and reduced reversibility, all consistent with biofouling from biomolecule adsorption. EIS in DA confirmed a marked increase in R_CT_ after incubation (**Figure S10D**). SEM revealed no morphological degradation (**Figure S10E– F**), while EDX (**Figure S10G)** spectra showed non-significant variations in Au/C ratio (**Figure S10H**), excluding Au NPs loss. The concurrent increase in the N/C ratio instead indicated adsorption of nitrogen-rich biomolecules from the medium. Together, these results confirm that the 20% signal loss observed in the long-term stability test arises from mild biofouling caused by prolonged exposure to cell culture medium. Overall, these results demonstrate that the sensor is only partially affected by biofouling and retains sufficient residual sensitivity to monitor neurotransmitter concentrations in neuronal cultures.

### 2.2. Differentiation of iPSCs into DAergic and GLUTergic neuronal cultures

To move beyond conventional *in vitro* sensing platforms, relying on immortalized cell lines, we implemented a human iPSC-derived neuronal model directly interfaced with the electroactive substrate. This approach enables the evaluation of electrochemical sensing performance within a physiologically relevant DAergic system, capturing the complex interplay between DA synthesis, release, and reuptake.

The platform’s performance was therefore validated using human DAergic neurons, which provide a more physiologically faithful model than traditional immortalized cell lines. By recapitulating the complex release and uptake dynamics of DA, these neurons enable a rigorous evaluation of both electrochemical sensitivity and long-term biocompatibility of the platform in a context that closely mimics the human brain environment.

In this work, we optimized existing differentiation protocols^[47]^ to differentiate iPSC into both DAergic and GLUTergic neurons. Furthermore, we investigated the effect of CNTs on subtype-specific differentiation, gene expression, and functional activity.

To evaluate the effect of CNTs on cellular differentiation, we modified the culture support by drop casting the bottom of the wells with the nanomaterial. Cells were cultured in DAergic neuron conversion medium until Day 13, with the medium refreshed every two days.

On Day 13, the medium was switched to a neuron maturation medium. From Day 13 to Day 24-30, the medium was refreshed every two or three days. The full composition of both media is detailed in the Supporting Information (SI). **Figure 2A** presents optical micrographs of DAergic neurons, illustrating the morphological changes at various stages of this process. On Day 0, the cells are observed as densely packed colonies with a uniform, compact morphology, a hallmark of undifferentiated iPSCs. By Day 18, the cells begin to undergo significant morphological changes, adopting a more neuron-like appearance with the emergence of projections. By Days 22-25, the cells display a mature neuronal morphology, characterized by an extensive, interconnected network of projections and small cell bodies (somas), indicating advanced neuronal differentiation. This progressive development highlights the successful transition from pluripotent stem cells to fully differentiated neurons. Further analysis was conducted on Day 28 using confocal microscopy and immunostaining to assess differentiation and morphological phenotype. In both the control and CNT groups, the DAergic neurons expressed neurofilament, confirming their neuronal identity (**Figure 2B**). To specifically confirm their DAergic phenotype, tyrosine hydroxylase (TH), a key enzyme in DA biosynthesis, was used as a marker. The neurons showed TH immunopositivity, validating their DAergic identity. Quantitative analysis revealed that the presence of CNTs significantly increased the efficiency of DAergic differentiation, defined here as the fraction of the neurofilament-positive area that was also TH-positive. The CNT group had a TH/neurofilament expression ratio of 37.90 ± 3.80%, which was significantly higher than the control group’s ratio of 26.01 ± 3.75% (p < 0.05) (**Figure 2C**).

**Figure 2.**
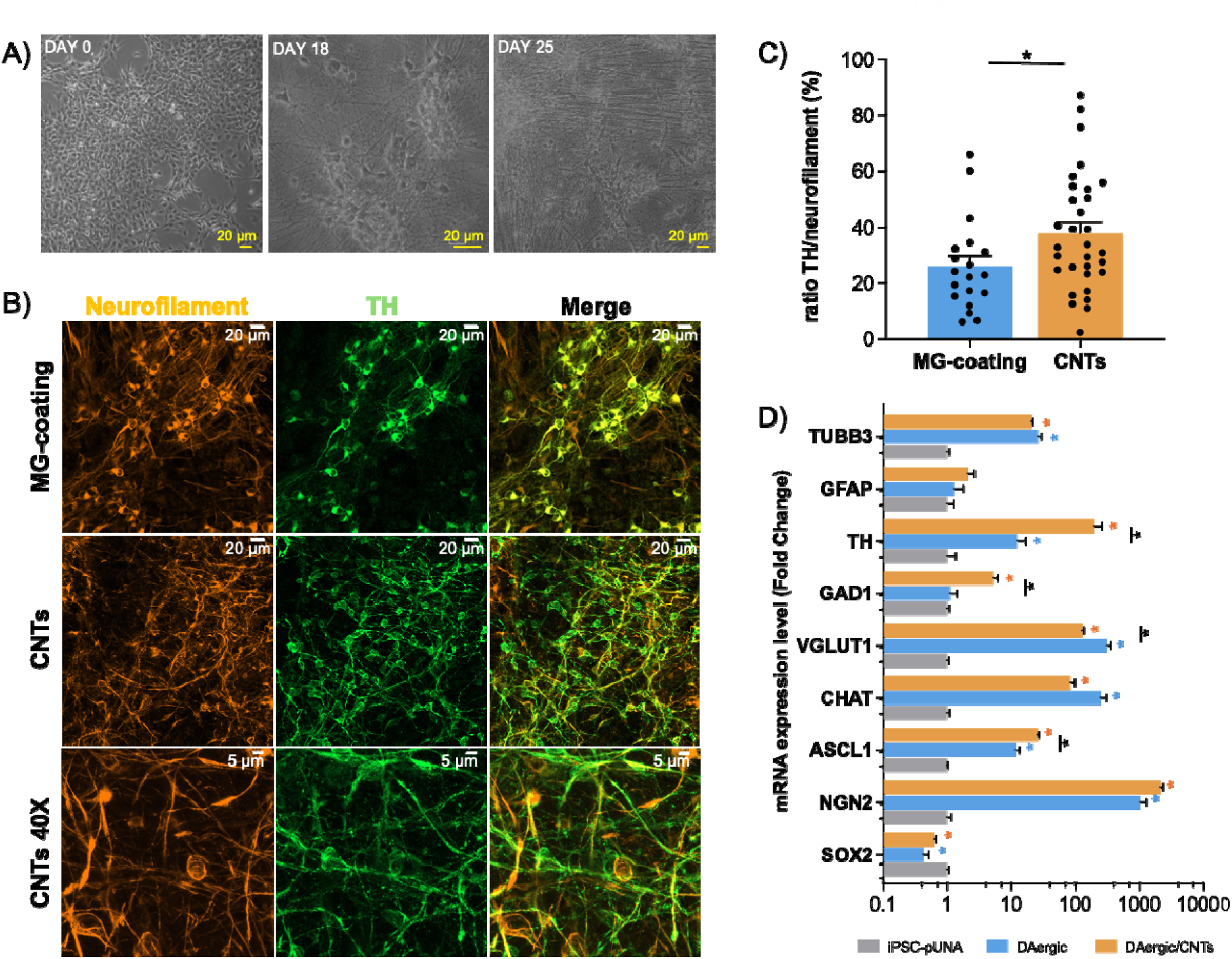
Morphological and phenotypical characterization of iPSC-derived DAergic neurons on CNTs. A) Optical micrographs of DAergic neurons at different days of differentiation. B) Immunostaining micrographs of DAergic neurons: control at 20X, in the presence of CNTs at 20X, and CNTs at 40X. iPSCs undergo differentiation, and after 28 days, the neurons were stained for neurofilament and TH. The merged staining is also shown. C) Quantification of the % overlap between neurofilament and TH in the images was performed (mean ± SEOM), *p < 0.05. The data were normally distributed, and a t-test was used to analyze differences between samples. D) Cellular characterization of DAergic neurons by quantitative qPCR. The levels of mRNA expression were measured by qPCR and calculated relative to the geometric mean of GAPDH-mRNA and Actin-mRNA reference controls using the 2^-ΔΔCt^ method. Each column represents the mean of three biological and technical replicates. The bars represent the mean ± SEOM (n=4). Statistical differences were analyzed between the control and the DAergic neurons with and without CNTs using the Mann-Whitney test. mRNA expression levels of SOX2, Ascl1, Ngn2, TUBB3, TH, GAD1, vGLUT1, CHAT and GFAP. Blue asterisks denote comparisons between iPSC-pUNA and DAergic neurons, orange markers highlight comparisons between iPSC-pUNA and DAergic/CNTs, and black markers denote differences between iPSC-pUNA and DAergic/CNTs.

To assess the gene expression profiles of the neuronal cultures, we performed quantitative Polymerase Chain Reaction (qPCR). We examined various culture conditions: undifferentiated iPSC-pUNA as a control, and DAergic neurons both with and without exposure to CNTs (**Figure 2D**). Our analysis confirmed the loss of pluripotency, as shown by decreased SOX2 levels in both differentiation conditions, indicating the absence of residual iPSCs. Quantification of NGN2 and ASCL1 mRNA levels revealed a significant increase in both genes’ expression in differentiated neurons compared to the iPSC-pUNA control, which confirmed successful induction of expression. Furthermore, the statistically significant increase in ASCL1 levels with CNTs suggests that this material may enhance the efficacy of the DAergic neuronal induction protocol, potentially increasing the proportion of cells competent to differentiate into neurons. The presence of TUBB3, a pan-neuronal marker, in both DAergic neuron groups validated the successful differentiation into neuronal cultures. We further characterized the neuronal composition by analyzing subtype-specific markers. Our results confirm that the induction protocol yields a mixed neuronal population rather than a pure culture, featuring GLUTergic (vGLUT1), GABAergic (GAD1), and cholinergic (ChAT) subpopulations alongside the enriched DAergic (TH+) neurons. Rather than a lack of specificity, this cellular diversity provides a multi-transmitter network that more closely mirrors the physiological complexity of the human midbrain. Notably, the presence of excitatory neurons, such as the GLUTergic population, may synergistically support the functional maturation and DA release dynamics within the culture. Interestingly, the presence of CNTs was associated with a change in this distribution. While the overall neuronal content remained comparable, as indicated by stable TUBB3 levels, CNT-exposed cultures showed a profile enriched in DAergic and GABAergic markers, with a marked increase in TH and GAD1 expression. Conversely, GLUTergic (vGLUT1) and cholinergic (ChAT) markers were downregulated in the presence of the nanotubes. These findings suggest that CNTs do not merely act as a passive structural scaffold but influence the subtype composition of the culture, favoring a DAergic profile over other subtypes. Whether this reflects altered fate specification, preferential maturation or survival of specific subpopulations, or changes in the expression of subtype-associated genes cannot be discriminated with the present data. Nevertheless, the reciprocal changes in subtype markers, together with unchanged TUBB3 expression, argue against a purely global effect on neuronal maturation.

To further contextualize the maturation of our model, we compared the DAergic induction protocol with a GLUTergic differentiation pipeline. As shown in **Figure 3A**, GLUTergic neurons exhibited significantly faster maturation kinetics compared to their DAergic counterparts. While DAergic cultures required a more extended period to establish complex structures, initial neuronal morphologies in GLUTergic cultures were clearly discernible by day 6. By day 18, the GLUTergic networks reached an advanced developmental stage, characterized by the clustering of neuronal somas and the formation of dense, parallel axonal projections.

**Figure 3.**
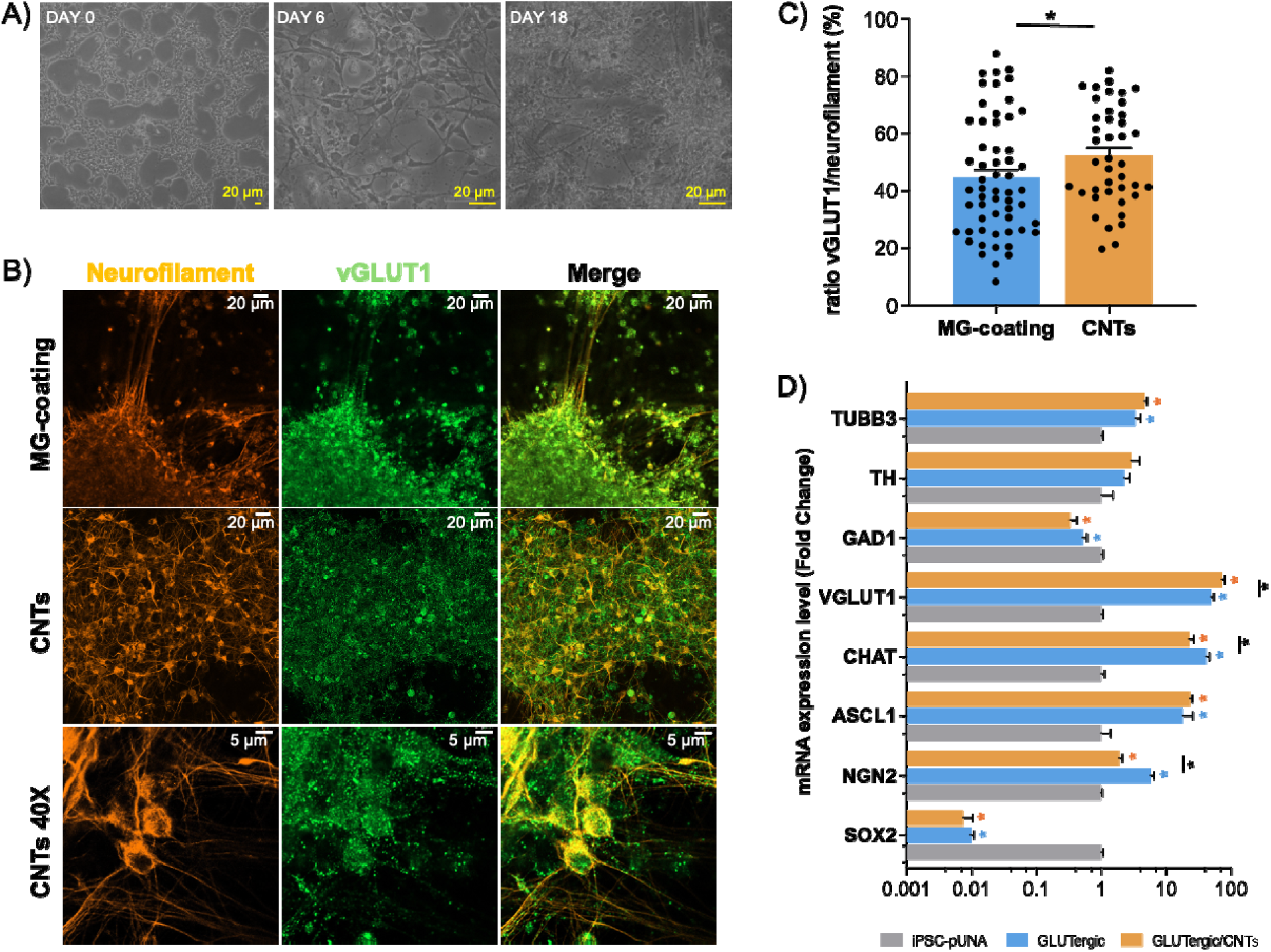
Morphological and phenotypical characterization of iPSC-derived GLUTergic neurons on CNTs. A) Optical micrographs of GLUTergic neurons at different days of differentiation. B) Immunostaining micrographs of GLUTergic neurons: control at 20X, in the presence of CNTs at 20X, and CNTs at 40X. iPSCs underwent differentiation, and after 24 days, the neurons were stained for neurofilament and vGLUT1. The merged staining is also shown. C) Quantification of the % overlap between neurofilament and vGLUT1 of the images was performed (mean ± SEOM), * p < 0.05. The data were normally distributed, and a t-test was used to analyze differences between samples. D) Cellular characterization of GLUTergic neurons by quantitative qPCR. The levels of mRNA expression were measured by qPCR and calculated relative to the geometric mean of GAPDH-mRNA and Actin-mRNA reference controls using the 2^-ΔΔCt^ method. Each column represents the mean of three biological and technical replicates. The bars represent the mean ± SEOM (n=4). Statistical differences were analyzed between the control and the GLUTergic neurons with and without CNTs using the Mann-Whitney test. mRNA expression levels of SOX2, Ascl1, Ngn2, TUBB3, TH, GAD1, vGLUT1 and CHAT were analyzed. Blue asterisks denote comparisons between iPSC-pUNA and GLUTergic neurons, orange markers highlight comparisons between iPSC-pUNA and GLUTergic /CNTs, and black markers denote differences between GLUTergic neurons with or without CNTs.

This rapid development led to a highly interconnected network earlier in the differentiation process, highlighting the distinct temporal dynamics inherent to each neuronal lineage.

Immunostaining showed that the GLUTergic neurons were actually positive for vGLUT1, with a punctate cytoplasmic expression that confirmed the GLUTergic phenotype (**Figure 3B**). We quantified the efficiency of GLUTergic differentiation by calculating the percentage of the neurofilament-positive area that was also vGLUT1-positive. This analysis revealed a significant increase in the presence of CNT substrates (overlap of 52.40 ± 2.72%), compared to the control group (44.63 ± 2.73%; p < 0.05) (**Figure 3C**). These results suggest that CNTs are also associated with enhanced GLUTergic differentiation.

The qPCR analysis for the GLUTergic protocol (**Figure 3D**) showed a gene expression profile consistent with our previous observations in DAergic cultures. Specifically, the downregulation of SOX2 alongside the robust activation of NGN2, ASCL1, and TUBB3 confirmed the successful exit from pluripotency and the acquisition of a neuronal identity. However, differentiation specificity was confirmed by a significant increase in vGLUT1 mRNA. The absence of spontaneous TH or GAD1 expression in these cultures highlights that the specific DAergic induction environment is essential to drive the DAergic phenotype.

The presence of CNTs further elevated vGLUT1 expression, suggesting that CNTs promote GLUTergic differentiation or enhance the expression of the glutamatergic vesicular machinery. Interestingly, ChAT expression was reduced in the presence of CNTs. Together with the observations in DAergic cultures, these results indicate that CNTs do not impose a fixed neuronal identity but rather reinforce the subtype directed by each induction protocol.

Although a definitive mechanistic explanation is beyond the scope of this study, a plausible framework can be proposed for our observation, based on previous reports. CNT nano-topography has been shown to activate integrin-mediated focal adhesion signaling and FAK phosphorylation, thereby influencing neuronal differentiation.^[32]^ Downstream activation of pathways such as FAK–AKT–β-catenin is particularly relevant, as canonical Wnt/β-catenin signaling is a well-established driver of DAergic specification and suppressor of alternative neuronal lineages.^[48–50]^ Additionally, the electrical properties of CNTs may modulate intracellular calcium dynamics, a known regulator of neurotransmitter subtype specification.^[51]^ These previously described pathways may contribute to the observed enrichment of specific neuronal subtypes; however, they were not directly assessed in this study and should be regarded as hypotheses. Their direct validation, for example through pharmacological inhibition of FAK or Wnt/β-catenin signaling during differentiation on CNT substrates, will be required to establish the underlying mechanism.

Following phenotypic and genotypic validation of neuronal cultures, we performed functional phenotyping experiments using a MEA system. This non-invasive platform allows for the longitudinal monitoring of emergent network behavior through three core electrophysiological metrics. The mean firing rate (MFR) provides a baseline of overall network excitability and tonic activity. The burst percentage (BP), the proportion of action potentials occurring within high-frequency clusters, serves as a robust indicator of synaptic strength and the refinement of neuronal signaling. Finally, network burst frequency (NBF) quantifies the occurrence of synchronized firing events across multiple electrodes, a hallmark of mature, integrated circuit formation. Collectively, these metrics offer a comprehensive picture of the functional development of the neuronal network.

Both DAergic and GLUTergic populations exhibited robust spontaneous electrical activity, confirming the functional maturation of the iPSC-derived networks. In DAergic cultures (**Figure 4A-C**), the MFR and burst percentage remained remarkably stable throughout the 55-day recording period, indicating a steady state of neuronal excitability and signaling. Although the network burst frequency showed an initial decline, it stabilized after day 30, maintaining values consistent with a functionally mature and integrated DAergic network. In contrast, GLUTergic neurons (**Figure 4D-F**) displayed more dynamic electrophysiological profiles. Following day 30, the MFR increased significantly, reaching peak activity between days 40 and 50 before stabilizing at later stages. This was accompanied by a rapid rise in burst percentage, which plateaued at high levels (75–85%) between days 30 and 60, reflecting a high degree of synaptic efficiency. The network burst frequency exhibited a pronounced peak around day 40, followed by a sharp decline, suggesting a period of intense network remodeling and synchronized firing during mid-differentiation. Collectively, these data demonstrate that both lineages achieve functional competence from day 24 onward. While GLUTergic networks exhibit a more transient peak in synchronized connectivity, both models maintain mature individual firing patterns and sustained burst activity through the later stages of culture.

**Figure 4.**
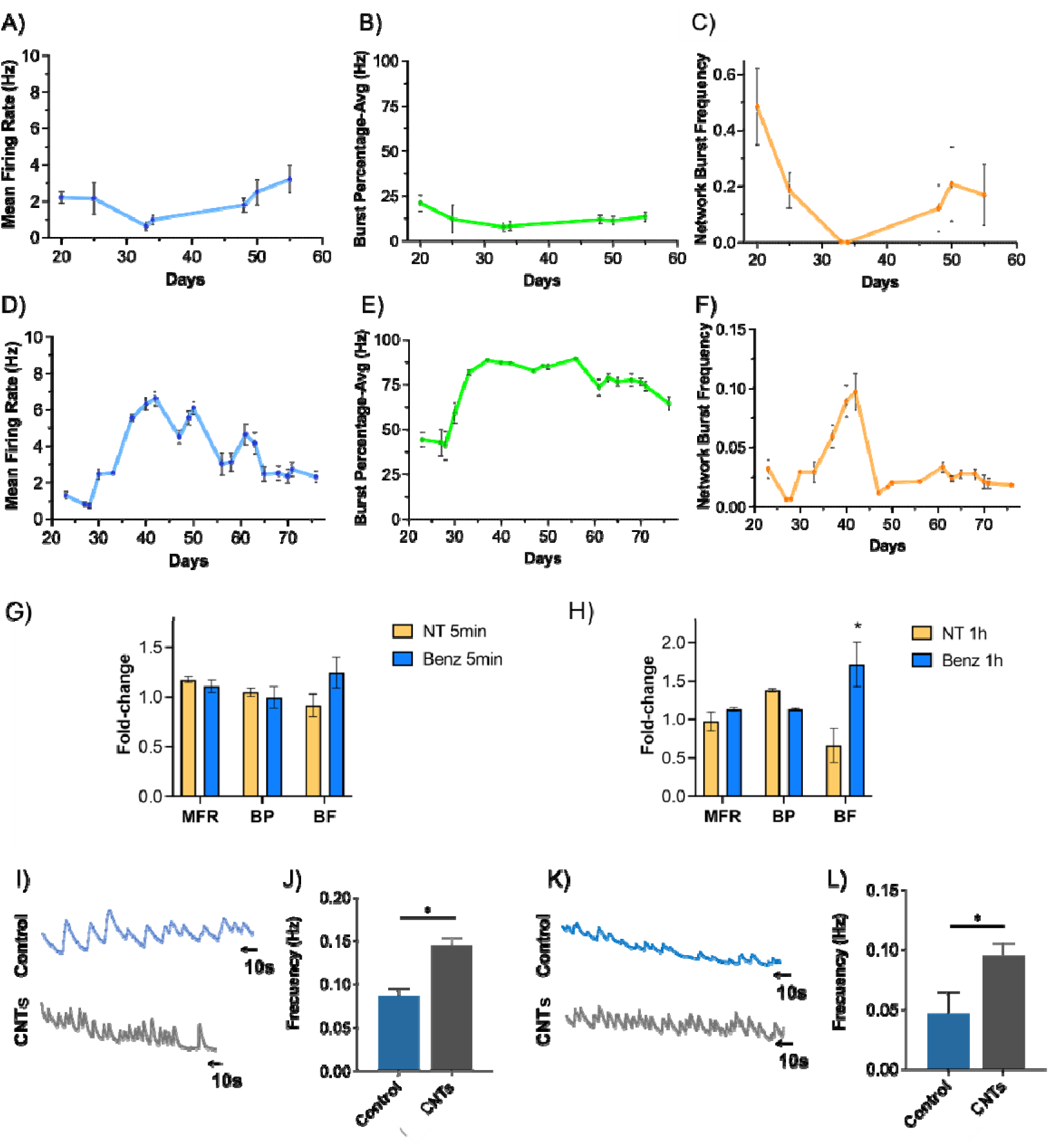
Long-term functional characterization of iPSC-derived neuronal networks via MEA electrophysiology and calcium imaging. A–C) Electrophysiological profile of DAergic cultures. Longitudinal monitoring from day in vitro (DIV) 21 to 55 (n=12) showing: A) Mean Firing Rate (MFR), B) Burst Percentage (BP), and C) Network Burst Frequency (NBF). D–F) Electrophysiological profile of GLUTergic cultures. Monitoring from DIV 21 to 76 (n=12) showing: D) MFR, E) BP, and F) NBF. All MEA data are presented as mean ± SEOM. G–H) Pharmacological modulation of DAergic activity. Effect of benztropine (DAT inhibitor) on MFR, BP, and NBF at G) 5 minutes and H) 1 hour post-treatment. Data are expressed as fold-change relative to non-treated controls and presented as mean ± SEOM from n = 3 independent biological replicates (individual differentiation wells). Statistical significance was determined using Student’s paired t-test; p < 0.05. I–L) Analysis of spontaneous neuronal calcium dynamics. I) Representative calcium transient plots and J) quantitative comparison of signal frequency between control and CNT groups in DAergic neurons. K) Representative calcium transient plots and L) frequency comparison in GLUTergic neurons. Data are represented as mean ± SEOM.Statistical significance was determined using Student’s paired t-test; p < 0.05.

To validate the functional DA machinery, we examined the response of DAergic cultures to benztropine, a selective DA transporter (DAT) inhibitor (**Figure 4G-H**). While no immediate changes were observed after 5 minutes, likely due to inhibitory effects through D2 autoreceptors, 1 hour of treatment induced a noticeable, significant increase in NBF (p < 0.05). This delayed response suggests that sustained extracellular DA accumulation modulates network excitability, potentially through the activation of excitatory postsynaptic receptors or complex network-level synchronization. These findings confirm the presence of active DA release and reuptake mechanisms, validating the physiological relevance of our iPSC-derived model. However, as MEA-derived metrics primarily reflect slow, adaptive network shifts in response to pharmacological modulation, additional methods with higher temporal resolution are required to directly quantify synaptic DA release and its immediate oxidative kinetics.

Calcium imaging was employed as a complementary approach to evaluate the impact of CNTs on neuronal functional activity, as the high electrical conductivity of these materials precludes their use in MEA systems due to the risk of inter-electrode short-circuiting. Spontaneous calcium transients were recorded from DAergic and GLUTergic neurons at day 26, both with and without CNTs. In both neuron types, spontaneous calcium oscillations were observed in the absence of external electrostimulation, further confirming their functional maturation. Quantitative analysis revealed that control DAergic neurons cultured on CNT-modified surfaces exhibited a significantly higher transient frequency (0.145 ± 0.009 Hz) compared to controls (0.086 ± 0.008 Hz; **Figure 4I-J**). A similar trend was found in GLUTergic neurons, where the presence of CNTs more than doubled the frequency of calcium events (0.095 ± 0.01 Hz vs. 0.046 ± 0.01 Hz in controls; **Figure 4K-L**). These data indicate that spontaneous neuronal activity is higher in the presence of CNTs, which may contribute to increased neurotransmitter release. Consequently, this CNT-augmented model provides a robust biological environment for validating the performance of our electroactive cell-culturing platform.

Beyond serving as a sensing interface, the CNT-based platform revealed an unexpected biological effect: an enrichment of the protocol-directed neuronal subtype during iPSC differentiation. This effect was consistently observed across immunocytochemical and transcriptional analyses, suggesting that CNTs influence the outcome of differentiation rather than acting solely as a passive structural scaffold. These findings position the material not only as an electrochemical enhancer but also as a substrate that promotes subtype-specific differentiation in human-derived systems. This observation is particularly relevant in light of previous reports, where CNT-based systems have been shown to promote general neuronal differentiation and maturation but have not demonstrated a clear effect on subtype-specific differentiation. In most cases, DAergic features are inferred from model systems with pre-existing phenotypic bias, rather than induced de novo. Our results, therefore, suggest that CNT substrates may influence neuronal subtype composition to a greater extent than previously recognized, although the underlying molecular mechanism remains to be established.

### 2.3. DA measurement in cellular cultures

In summary, our protocol generates TH-positive neurons that exhibit robust spontaneous electrophysiological activity and a functional DA reuptake machinery, as confirmed by MEA and pharmacological assays. This human-based model reproduces key electrophysiological and neurochemical features of DAergic neurons, including spontaneous network activity and functional DA reuptake. Consequently, it provides a suitable biological model to validate our electroactive platform for the *in situ* detection of DA.

Before beginning the in-vitro experiments, the biocompatibility of the Au-CNTs and CNTs materials was evaluated. A Total Cell Number Assay, which quantifies the release of lactate dehydrogenase (LDH) as a marker for cytotoxicity, was performed to assess their effect on cell viability.

For the assay, 100,000 iPSCs were seeded into 8-well chamber plates coated in quadruplicate with either CNTs or Au-CNTs at a concentration of 1 mg mL^-1^. Geltrex-coated wells were used as the control. After incubation, cell viability was determined through a colorimetric analysis. The results showed that both the CNT and Au-CNTs coatings exhibited good cell viability, comparable to the Geltrex control. (**Figure S11**) This confirmed that the materials were non-toxic and suitable for subsequent experiments.

To detect DA release at the cellular level, we differentiated iPSCs directly on the biosensor. We used five replicates for DAergic neurons and four for GLUTergic neurons.

Before the electrochemical measurements, we replaced the cell culture medium with Krebs-Ringer medium. We then performed DPV using optimized parameters. The first sweep of the DPV did not show any significant oxidation peak for DA. However, a well-defined peak appeared at 0.27 V starting from the second sweep, which we correlated with measurement-induced release of DA from the neurons. The peak current intensity increased with each sweep until the third sweep, after which it stabilized (**Figure 5A**). This result suggests that the sweep applied during DPV produces a localized interfacial current density at the cell–electrode interface that is sufficient to depolarize the neuronal membrane and trigger exocytotic DA release. Across the five replicates, the average current measured at the third sweep was 7.27 ± 3.48 µA (**Figure 5B**). The high standard deviation is likely due to the inherent variability of biological differentiation processes. Based on the previously reported observations, our interpretation is that this current value is related to measurement-induced DA concentration, rather than a basal extracellular concentration.

**Figure 5.**
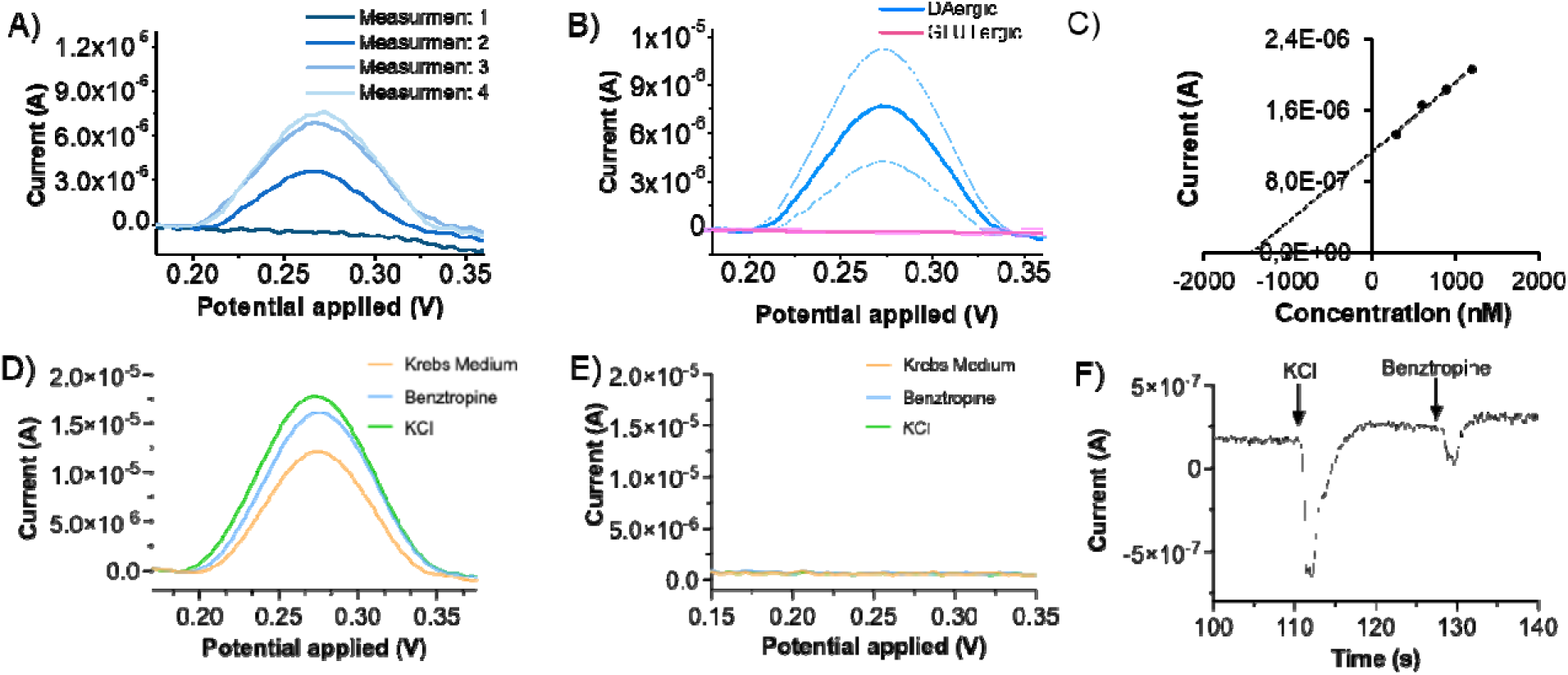
A) DPV response registered from DAergic neurons in Krebs-Ringer medium after consecutive measurements. B) Comparison of the average DPV response at the third measurement (n=5) from DAergic and GLUTergic neurons. C) Standard addition calibration curve for DA concentration in the range of 300 nM to 1.8 µM obtained in DAergic culture. D) DPV response registered from DAergic neurons before and after the addition of benztropine and KCl. E) DPV response registered from GLUTergic neurons before and after the addition of benztropine and KCl. F) Chronoamperogram of DAergic neurons upon addition of KCl and benztropine.

GLUTergic neurons, which are not known to produce DA, were employed as a negative control. In the same experimental conditions, these neurons did not show any faradaic current even after the third DPV sweep (**Figure 5B**). This result indicates that the appearance of the redox signal requires the presence of DAergic cells and is not attributable to nonspecific electrode effects, general cell–surface interactions, or oxidation of medium components.

To determine the concentration of DA released by DAergic neurons after the measurement-induced membrane polarization, we employed the standard addition method. This internal calibration strategy, in which known concentrations of DA are directly spiked into the cell culture, effectively corrects for both matrix-induced signal attenuation and the sensitivity changes over time observed in the long-term stability tests (**Figure 1H**), ensuring accurate and reliable DA quantification.

Initially, we performed DPV sweeps to stimulate measurements-induced release of DA, until a stable DA oxidation signal was achieved. Subsequently, DA standard concentrations were added to the cell culture, and the corresponding current was measured via DPV. The DA concentration was extrapolated as the absolute value of the intercept of the standard calibration line with the x-axis (**Figure 5C**). The calculated value of 1.25 µM is higher than the endogenous levels typically observed in PC12 cell cultures (**Table S11**). This elevated concentration can be attributed to three possible and concomitant explanations: 1) it represents the DPV-induced DA concentration, rather than a basal extracellular concentration; 2) it reflects the high functionality, spike frequency, and elevated DA production of the cell culture itself; 3) the surface-sensitive nature of electrochemical techniques probes a localized diffusion layer within a few micrometers of the electrode surface. Therefore, although the spatial origin of the signal remains uninvestigated in the present study, we hypothesize that the sensor detects DA released locally by neurons growing directly on the sensing surface.

After demonstrating that the electroactive platforms could detect and quantify DA released by neurons, we investigated their ability to measure changes in DA levels induced by drugs or external stimuli. We treated three out of five electrodes, which were cultured with DAergic neurons, with benztropine. This Parkinson’s disease medication blocks DA reuptake, thereby increasing its availability in the brain. After the measurement-induced DA levels were stabilized, we added benztropine to the culture to achieve a final concentration of 400 nM. The subsequent inhibition of DA reuptake resulted in a prolonged presence of extracellular DA, which was reflected by a significant increase in the current (**Figure 5D**). Although the initial basal currents varied among the three electrodes, the benztropine treatment consistently increased the current due to DA oxidation by 43.59 ± 10.77% (**Figure S12**). This confirms that our platform specifically responds to DA release from DAergic neurons and can be used to assess drug efficacy.

A further stimulus was applied by incubating the cell culture with 80 mM KCl. Potassium ions (K^+^) induce neuronal membrane depolarization, which stimulates the release of DA from the synapses. This also resulted in a consistent current increase of 17.08 ± 3.92%, confirming a rise in DA concentration.

On the other hand, the GLUTergic neurons never released DA in detectable amounts, nor after the treatment with benztropine or KCl (**Figure 5E**), as expected.

A complementary experiment using chronoamperometry (CA) was conducted to further validate our findings (**Figure 5F**). By applying a constant potential of 0.27 V, we initially observed a flat and stable baseline current, indicating that this potential was not high enough to depolarize the neuronal membrane and trigger exocytotic DA release.

Following the addition of KCl and benztropine to the cell culture, a distinct two-step increase in current was observed. The sharp spikes seen during the additions are attributed to turbulent mixing and should not be considered. Cell-free control experiments (**Figure S13**) confirmed that adding KCl to 1.5 µM DA produced no current change in either DPV or CA, confirming that the current responses observed in neuronal cell cultures originate specifically from depolarization-induced DA exocytosis rather than non-specific changes in ionic strength or solution conductivity

This experiment confirms the specific response of the neurons to these chemical stimuli, proving that DA levels can also be monitored via CA. The results also validate that the current increases observed during the DPV experiments were indeed due to the compounds introduced, and not merely a consequence of the electrical stimulation.

After the DPV and CA measurements, immunostaining was performed on the DAergic and GLUTergic cell cultures to confirm neuronal integrity and differentiation. (**Figure 6**) This post-experimental validation was essential to verify that the neurons retained their morphology and marker expression after the electrochemical testing. For the immunostaining, neurofilament, TH, and DAPI were used as markers for both types of neurons. As expected, the DAergic neurons were found to be mature and TH-positive (TH+), confirming their DAergic identity. The GLUTergic neurons only exhibited positive staining for neurofilament and DAPI but were negative for TH, confirming their role as a negative control for DA release. This validation step also confirmed that the voltage applied during the electrochemical measurements had no evident adverse effects on neuronal morphology or marker expression.

**Figure 6.**
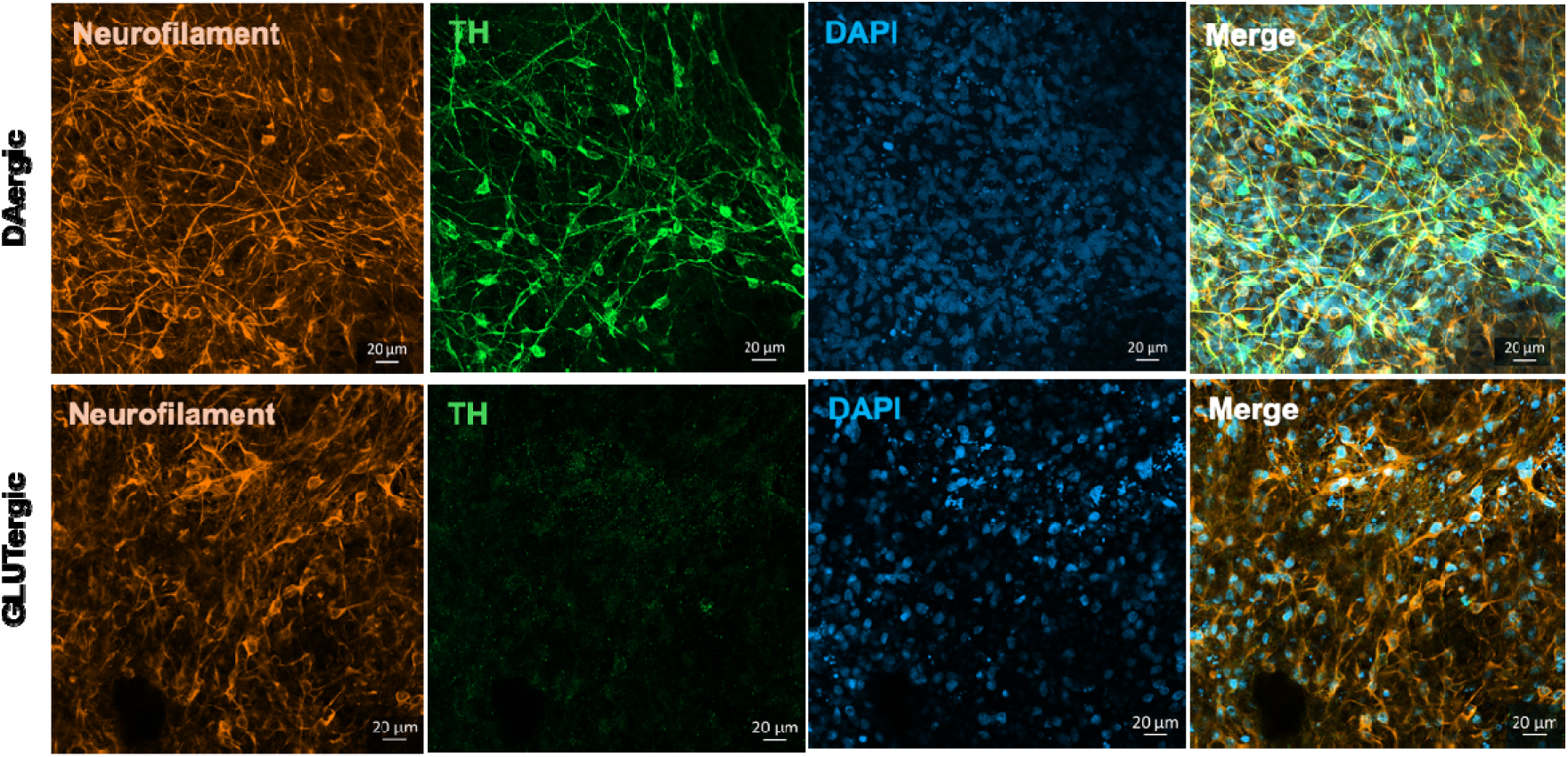
Composite fluorescence images of immunostained DAergic and GLUTergic neurons on ITO modified with CNT/Au-CNT at day 26. The samples were stained for neurofilament (Orange), TH (green), DAPI (blue), and the merge is also shown; magnification 20x. Scale bar: 20 μm.

## 3. Conclusion

In conclusion, this study presents an electrochemical neurotransmitter sensing platform that integrates CNT-modified electrochemical sensors with a physiologically relevant human DAergic neuronal model. Compared to widely used systems based on immortalized cell lines (e.g., PC12) or acellular calibration assays, the use of iPSC-derived human neurons provides a more physiologically relevant context to study DA synthesis, release, and uptake. In line with previous reports employing CNT-modified electrodes for DA detection, our platform confirms the well-known gains in electrochemical sensitivity and selectivity; however, it goes beyond prior studies by demonstrating long-term biocompatibility, enhanced subtype-specific differentiation and increased spontaneous neuronal activity directly on the sensing interface. The multiple role of CNTs as electrocatalytic enhancers and substrates that promote subtype-specific differentiation represents a clear advance over passive electrode coatings reported in most brain-on-a-chip and neurobiosensing studies. Importantly, the platform detects cell-derived DA and its pharmacological modulation *in situ*, a capability that remains largely inaccessible to conventional HPLC-, ELISA-, or MEA-based approaches. These features support the potential of the system as a scalable and versatile phenotypic screening platform for DAergic modulators, with direct relevance to Parkinson’s disease research. While CNT-based electrodes, iPSC-derived neurons, and electrochemical DA sensing have each been reported separately, this work uniquely integrates all three in a single platform where CNTs simultaneously promote subtype-specific differentiation and enhance analytical sensitivity More broadly, the modular design and human-cell-based validation suggest strong potential for extension to other neurotransmitter systems and neurological disorders, supporting its future application in drug discovery, disease modeling, and precision neuropharmacology. Importantly, this work expands the role of CNT-based materials beyond that of passive electrochemical modifiers, showing that they can also promote subtype-specific differentiation and enhance the spontaneous activity of human iPSC-derived neurons.

## 4. Experimental section

### iPSC culture and viral transfection

Human induced pluripotent stem cells (iPSCs, JIPSC1100) were acquired from The Jackson Laboratory. The iPSCs were cultured in mTeSR™ Plus (Stem Cell) supplemented with penicillin/streptomycin (100 mg mL^−1^Gibco). Cultures were maintained at 37°C in a humidified atmosphere with 5% CO_2_. For routine maintenance, culture plates were coated overnight with Corning™ Matrigel™ membrane matrix (Fisher), diluted 1:40 in Iscove’s Modified Dulbebe’s Medium (IMDM). T25 and T75 culture flasks were seeded with 1×10^6^ and 2.5×10^6^ cells, respectively. To pass the cells, the spent medium was removed, and the cells were incubated with Accutase™ for 5 minutes at 37°C. The detached cells were centrifuged at 300×g for 5 minutes, resuspended in mTeSR™ Plus supplemented with 1:1000 ROCK inhibitor, and reseeded. Passaging was performed regularly to prevent cell overgrowth and spontaneous differentiation.

iPSC-pUNA cells were generated by transducing iPSCs with lentiviral particles for the inducible expression of Ngn2 and Ascl1. These particles were produced using the pLVX-UbC-rtTA-Ngn2:2A:Ascl1 plasmid (Addgene, #127289), which allows for doxycycline-dependent transgene activation via the Tet-On system. Viral particles were produced by the Viral Vectors Unit of the Spanish National Center for Cardiovascular Research (CNIC), Madrid, Spain.

For transduction, iPSCs were seeded at 60% confluency in mTeSR™ Plus and treated with lentiviral particles at a multiplicity of infection (MOI) of 12. Polybrene was included to enhance cell membrane permeability. After 30 hours, the medium was replaced with fresh mTeSR™ Plus. Transduced cells were then selected using puromycin at a concentration of 0.35 μg mL^−1^, a concentration that was determined in preliminary experiments to kill all untransduced cells while maintaining the viability of transduced cells. Following selection, the cells were expanded in T25 flasks and maintained under puromycin selection in all subsequent passages.

### iPSC-pUNA differentiation into DAergic neurons

To obtain cultures enriched in DAergic neurons, we follow this procedure. First, puromycin-selected iPSCs-pUNA cells were cultured in various formats, including 8-well chambers, 6-well, 96-well, and Microelectrode Arrays (MEAs) plates (M384-tMEA-24W, Axion Biosystems). All culture surfaces were pre-coated with Geltrex™ LDEV-Free, an hESC-qualified, reduced growth factor basement membrane matrix (ThermoFisher, 12063569). A 1:100 dilution of Geltrex™ in IMDM was applied and incubated overnight at 37°C.

The following day, the Geltrex™ coating was removed, and iPSCs-pUNA cells were seeded into their respective formats. Cell densities varied by plate type: 500,000 cells for 6-well plates, 120,000 for 8-well chambers, 80,000 for 96-well plates, and 100,000 for MEAs (Day - 1). The seeding medium was supplemented with the ROCK inhibitor Y-27632 (StemCell Technologies) to target the RHO/ROCK pathway (ROCK1 and ROCK2). For MEAs, cells were seeded in a minimal volume of medium (7 μL) directly onto the electrodes and were allowed to attach for two hours before the remaining medium was added. The ROCK inhibitor was removed on the following day.

On Day 0, the mTeSR™ medium was replaced with a DAergic neuron conversion medium (composition detailed in the Supporting Information, **Table S6**). Cultures were maintained until Day 13, with the medium refreshed every two days. At day 13, the medium was refreshed with neuron maturation medium (composition detailed in the Supporting Information, **Table S7**). From day 12 to 24-30, until maturation, the medium was refreshed every two to three days.

### pUNA differentiation into GLUTergic neurons

To obtain a culture enriched in GLUTergic neurons, a similar culturing procedure used for the DAergic induction was followed, with few variations. On Day 0, the mTeSR™ medium was replaced with a neuron conversion medium (composition detailed in the Supporting Information, **Table S8**) until Day 8. The medium was completely replaced on Day 2. On Days 3 and 5, the medium was refreshed, and 10 mM DAPT (Selleckchem) was added. On Day 6, the medium was refreshed again with the neuron conversion medium supplemented with 10 mM DAPT and 5 mM Cytosine β-D-arabinofuranoside (AraC) (Sigma). AraC was included to inhibit the proliferation of non-neuronal cells. Typically, a single day of AraC treatment was sufficient; however, if cultures on Day 7 still contained cells with non-neuronal morphology, an additional day of AraC treatment was performed. The medium was refreshed again on Day 7. On Day 8, the medium was replaced with a Neuron Maturation Medium (composition detailed in the Supporting Information, **Table S9**). From Day 10 to Day 19-24, the medium was refreshed every two to three days.

### Immunostaining of iPSC-derived neurons

Upon reaching maturation (Day 24-26), the differentiated and matured neurons were fixed in 4% paraformaldehyde (PFA) for 20 minutes, followed by blocking and permeabilization in a solution containing 5% donkey serum, 0.1% Triton X-100 in phosphate-buffered saline (PBS), 2% BSA, and 0.1% NaN_3_ for 1 hour at room temperature. Primary antibodies were incubated overnight at 4°C and included: purified anti-Neurofilament H (NF-H), nonphosphorylated antibody (#SMI-32P, BioLegend), and anti-TH antibody (AB152, Sigma-Aldrich) or vGLUT1 (135 511, Synaptic Systems). After three washes with PBS, the cells were incubated for 2 hours with secondary antibodies (AlexaFluor488 and AlexaFluor647, Life Technologies) and DAPI (1 μg mL^−1^, D1306, Thermo Scientific). The samples were then rinsed three more times with PBS. Fluorescence micrographs were obtained using a Zeiss LSM 900 KMAT confocal microscope with AXIO OBSERVER 7.

The resulting immunostaining micrographs were analyzed using ImageJ software. To assess the proportion of neuronal cells differentiated into specific subtypes, the neurofilament-positive area was quantified and compared to the area stained for phenotype-specific markers, such as TH or vGLUT1. The percentage of the specific marker relative to the neurofilament area was calculated. Statistical significance of the results was determined using R Studio.

### Quantitative polymerase chain reaction

Total RNA was extracted using the RNase Mini Kit with an accompanying RNase-free DNase kit (Qiagen). RNA concentration was measured using a NanoDrop spectrophotometer, and 1 μg of RNA was used as the input for reverse transcription (RT). A total of 20 μL of complementary DNA (cDNA) was obtained from the RNA samples via RT using the High-Capacity kit (Life Technologies, Carlsbad, CA, USA). The cDNA was then mixed with 80 μL of H_2_O to achieve a stock concentration of 10 ng μL^−1^.

qPCR was performed on a CFX384 Touch qPCR Detection System (Bio-Rad) using Power SYBR Green Master Mix (Thermo Fisher), 300 nmol L^−1^ of each primer pair, and 10 ng of cDNA. Ct values were obtained using the Bio-Rad CFX Maestro software to analyze the plate. Two controls were included: a reverse transcription control (RTC), which lacked the multiscribe reverse transcriptase during the RT step, and a no template control (NTC), which used H_2_O instead of the cDNA sample.

The genes analyzed were Ascl1, Ngn2, TUBB3, TH, GAD1, vGLUT1, CHAT, and GFAP. Glyceraldehyde-3-phosphate dehydrogenase (GAPDH) and Actin served as housekeeping genes. The primer sequences for qPCR are listed in **Table S10**. Relative quantification was performed using the 2^−ΔΔCt^ method, and values were normalized to the non-differentiated iPSC controls (iPSC-pUNA).

### MEAS recordings

To characterize the maturation of iPSC-derived neuronal networks, we performed non-invasive extracellular electrophysiological recordings. Data were acquired using an Axion Maestro Edge MEA system operated with AxIS software (Axion Biosystems). For each recording session, spontaneous neuronal activity was measured for a duration of 8– 10 minutes within a controlled environment maintained at 37 °C and 5% CO_2_.

Raw signals were digitized at a sampling frequency of 12.5 kHz per channel. To isolate spikes from background noise and slow local field potentials, a digital band-pass filter of 1–300 Hz was applied. The neuronal cultures were maintained on CytoView MEA 24-well plates, each equipped with 16 integrated PEDOT-coated recording electrodes. Spike detection was performed using an adaptive thresholding algorithm set at 6x the standard deviation of the electrode’s RMS noise. The key parameters used for quantitative analysis of the network activity included the mean firing rate (MFR), burst percentage, and network burst frequency, which were calculated to assess the functional connectivity and synchronized firing patterns of the maturing networks.

### Calcium imaging and analysis

To assess neuronal activity, calcium imaging was conducted on iPSC-derived neurons at 24-28 days in vitro. Neuronal cultures were loaded with the fluorescent calcium indicator Fluo-8® AM (AAT Bioquest) at a final concentration of 2.5 µM and incubated for 15 minutes at 37 °C in the dark. Time-lapse imaging of calcium dynamics was performed using an inverted Nikon ECLIPSE Ti-S / L100 fluorescence microscope with a FITC filter. The study analyzed four independent batches, with three wells per condition, 12 videos per batch, and 16-18 regions of interest (ROIs) per video. Videos were recorded for 3 minutes at a frame rate of 500 ms/frame. Subsequent analysis of calcium signal frequency was carried out in MATLAB. Statistical analyses were performed using GraphPad Prism 10.4. Outliers were first identified and removed using the ROUT method (Q = 0.5%). The distribution of the remaining data was evaluated for normality and log-normality. Based on these results, a parametric unpaired t-test or a non-parametric Mann-Whitney test was used for statistical comparisons.

### Purification of the CNT and preparation of Au-CNT

CNTs were purchased from Nanostructures & Amorphous Materials Inc. (Stock# 1237YJS; inner diameter: 5–10 nm; outer diameter: 20–30 nm; length: 0.5–2 µm). For purification, 100 mg of CNTs were suspended in 6 M HCl and sonicated for one hour, followed by overnight agitation. The mixture was then diluted in 1 L of Milli-Q water and filtered through 0.45 µm filters until a neutral pH was achieved. XPS confirmed the absence of metallic impurities, and elemental analysis showed the CNTs were composed of 96.80% carbon (C) and 3.20% oxygen (O). The CNTs were decorated with gold nanoparticles (Au NPs) following the procedure reported in Misia et al.^[36]^

### Manufacture of the electroactive cell culturing platform

Indium tin oxide (ITO)-coated glass slides with a thickness of 0.16 mm µM (SPI Supplies; Ref. #06485-AB) were cut to 0.75×1.5 cm using a diamond-tip cutter. Subsequently, 0.75×0.75 cm polylactic acid (PLA) square wells were 3D-printed with a filament 3D printer. The PLA wells were then secured to one end of the ITO surface with SYLGARD™184 silicone elastomer to define the cell culture and electroactive area (0.75 cm^2^). The final electrochemical cell was completed with a platinum (Pt) wire as a counter electrode (CE) and a miniaturized Ag/AgCl reference electrode (RE).

### Modification of the culturing platforms with CNTs

The culture surfaces were modified by drop-casting CNT-based materials. For immunostaining, a 1 mg mL^−1^ aqueous CNT solution was drop-cast at a volume of 140 µL into each well of an 8-well chamber slide. For quantitative qPCR experiments, 1.5 mL of a 1 mg mL^−1^ CNT solution was spray-coated into each well of a 6-well plate. For electroactive culture plates, a two-step drop-casting procedure was followed. First, 60 µL of a 1 mg mL^−1^ aqueous CNT suspension was drop-cast and dried. A second layer of 120 µL of a 0.5 mg mL^−1^ Au-CNT suspension was then carefully drop-cast onto the CNT layer. After coating, all plates were placed on a hot plate at 40 °C to ensure uniform adhesion and drying. Before cell seeding, the plates were rinsed twice with water and sterilized by exposure to UV light for one hour.

### Electrochemical Measurements

All electrochemical measurements were performed using an Autolab MSTAT204 potentiostat/galvanostat (Metrohm Autolab) controlled by Nova 2.1.6 software. All experiments were conducted at room temperature.

In DPV, MA and MT were optimized using a Design of Experiments (DoE) approach (for further information, see the Supporting Information). The optimized parameters used were 0.052 s for MT and 0.062V for MA. In DPV measurements, the baseline was subtracted from the obtained voltammograms using a moving average baseline model.

To perform in vitro electrochemical experiments on DAergic neurons and GLUTergic controls, the cell culture medium was first removed. The cells were then washed and incubated in a Krebs-Ringer Buffer for 45 minutes at 37 °C in a 5% CO_2_ atmosphere before measurements. The buffer contained 146 mM NaCl, 2.7 mM KCl, 2.1 mM MgCl_2_, 1.2 mM CaCl_2_, 10 mM glucose, 15 mM HEPES, and 6 mM sucrose.

DPV measurements were carried out using the parameters optimized via the DoE. DPV was repeated until a constant signal was achieved. After signal stabilization, benztropine was added to reach a final concentration of 400 nM, and a subsequent DPV measurement was recorded. Finally, KCl was introduced to achieve a final concentration of 80 mM KCl, and a final DPV measurement was performed. For all DPV measurements, background currents were corrected by applying a moving average baseline subtraction algorithm in the NOVA software (Metrohm Autolab). Peak currents were subsequently determined from the baseline-subtracted voltammograms. In the absence of a distinct oxidation peak at low concentrations, the signal was sampled at the fixed potential corresponding to the peak maximum observed at higher DA concentrations.

The limit of detection (LOD) was determined based on the minimum distinguishable signal (*S_LOD_*), calculated as:

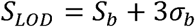

Where *S_b_* represents the mean background signal of the blank and *σ_b_* is the standard deviation of blank measurements across three independent electrodes. The concentration-based LOD was subsequently derived by substituting *S_LOD_* into the linear calibration equation (y = mx + q), yielding:

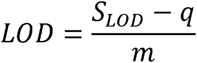

Where q and m correspond to the -intercept and slope of the calibration curve, respectively.

Chronoamperometry was performed at 0.16 V for 160 seconds. After the signal stabilized, KCl and benztropine were added sequentially while the signal continued to be recorded to reach final concentrations of 80 mM and 400 nM, respectively.

### Tafel analysis

The Tafel analysis was performed on the anodic (forward) branch of the cyclic voltammograms recorded with ITO/CNT/Au-CNT electrode at 5 mV s□¹ in the presence of 100 µM solution of each monoamine neurotransmitter over a potential range of -0.1 to 0.5 V. The current density (*j*) was determined on the geometric area of the ITO electrode (0.75 cm^2^). The Tafel slope was extracted from the linear region of the log |*j*| vs *E* plot in the rising portion of the anodic peak, before the onset of diffusion limitation.

### Electrochemical impedance spectroscopy

EIS measurements were acquired in Krebs-Ringer buffer solution containing 100 µM DA at a bias potential of E_1/2 =_ 0.18V vs. Ag/AgCl (corresponding to the formal half-wave potential of oxidation). A sinusoidal AC voltage perturbation of 10 mV amplitude was applied over a frequency range from 100 KHZ to 0.1 HZ . Impedance spectra were recorded across bare ITO, ITO/CNT, and double-layer ITO/CNT/Au-CNT electrodes.

### Long-Term Stability and Biofouling Evaluation

To evaluate fouling resistance and chronic electrochemical durability, ITO/CNT/Au-CNT electrodes were continuously incubated at 37 °C in a 1:1 mixture of DMEM:F12 and Neurobasal-Brainphys cellular growth medium for 21 days. To differentiate between biofouling-induced signal degradation and the intrinsic aging of the nanocomposite material, a parallel control group of electrodes was maintained under identical incubation conditions in a protein-free PBS solution. Every 3 days, the respective incubation matrices were temporarily removed, the electrodes were gently rinsed with deionized water, and DPV scans were performed in a fresh Krebs-Ringer buffer solution spiked with DA. All current responses were normalized to the baseline signal recorded on Day 0 to determine the percentage of signal change over the incubation time.

## Acknowledgements

A.S. acknowledges the financial support from Ca’ Foscari University of Venice.

## Author contributions

**Nerea Pascual Frías:** Conceptualization, Data curation, Formal analysis, Investigation, Methodology, Writing – original draft. **Giuseppe Misia**: Data curation, Formal analysis, Investigation, Methodology, Writing – review & editing; **Lara Rodríguez Sánchez**: Investigation, Formal analysis. **Claudio Evangelisti:** Investigation, Methodology, Writing – review & editing. **Maurizio Prato:** Funding acquisition, Resources, Supervision, Writing – review & editing; **Nuria Alegret:** Conceptualization, Funding acquisition, Methodology, Resources, Supervision, Writing – review & editing. **Francisco Gil Bea:** Conceptualization, Funding acquisition, Methodology, Resources, Supervision, Writing – review & editing. **Alessandro Silvestri:** Conceptualization, Data curation, Funding acquisition, Methodology, Investigation, Formal analysis, Resources, Supervision, Writing – original draft, Writing – review & editing.

Received: ((will be filled in by the editorial staff))

Revised: ((will be filled in by the editorial staff))

Published online: ((will be filled in by the editorial staff))

## Supporting Information

Supporting Information is available from the Wiley Online Library.

This work presents a novel CNT-based platform for dopamine quantification in living neurons. Carbon nanotubes act as conductive electrocatalysts and support neuronal growth, enabling DA detection in situ. Beyond sensing, CNT substrates promote iPSC differentiation into DAergic and GLUTergic neurons. The use of benztropine as a model drug demonstrated promising application in the field of neurobiological research and drug screening.

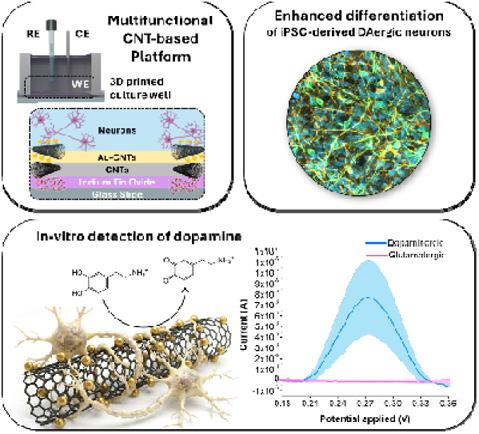

