## Supporting Information for "Multifunctional CNT-based Platform for Enhanced Dopaminergic Differentiation and Electrochemical Dopamine Sensing from Human iPSC-derived Neurons"

M. Prato

Department of Chemical and Pharmaceutical Sciences, INSTM Unit of Trieste, University of Trieste, Via L. Giorgieri 1, 34127 Trieste, Italy

M. Prato

Ikerbasque, Basque Foundation for Science, 48013 Bilbao, Spain

1. **Au-CNTs Characterization**

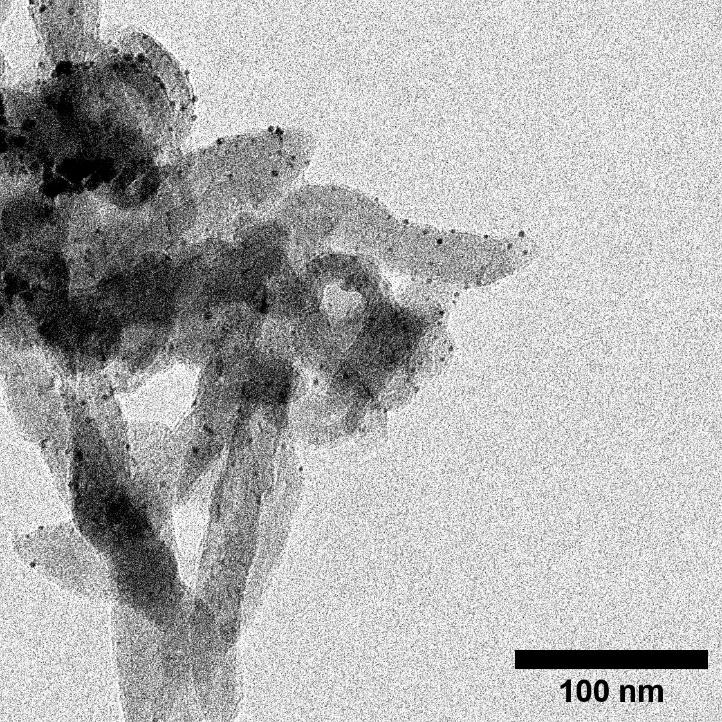

**Figure S1.** TEM micrograph of Au-CNTs.

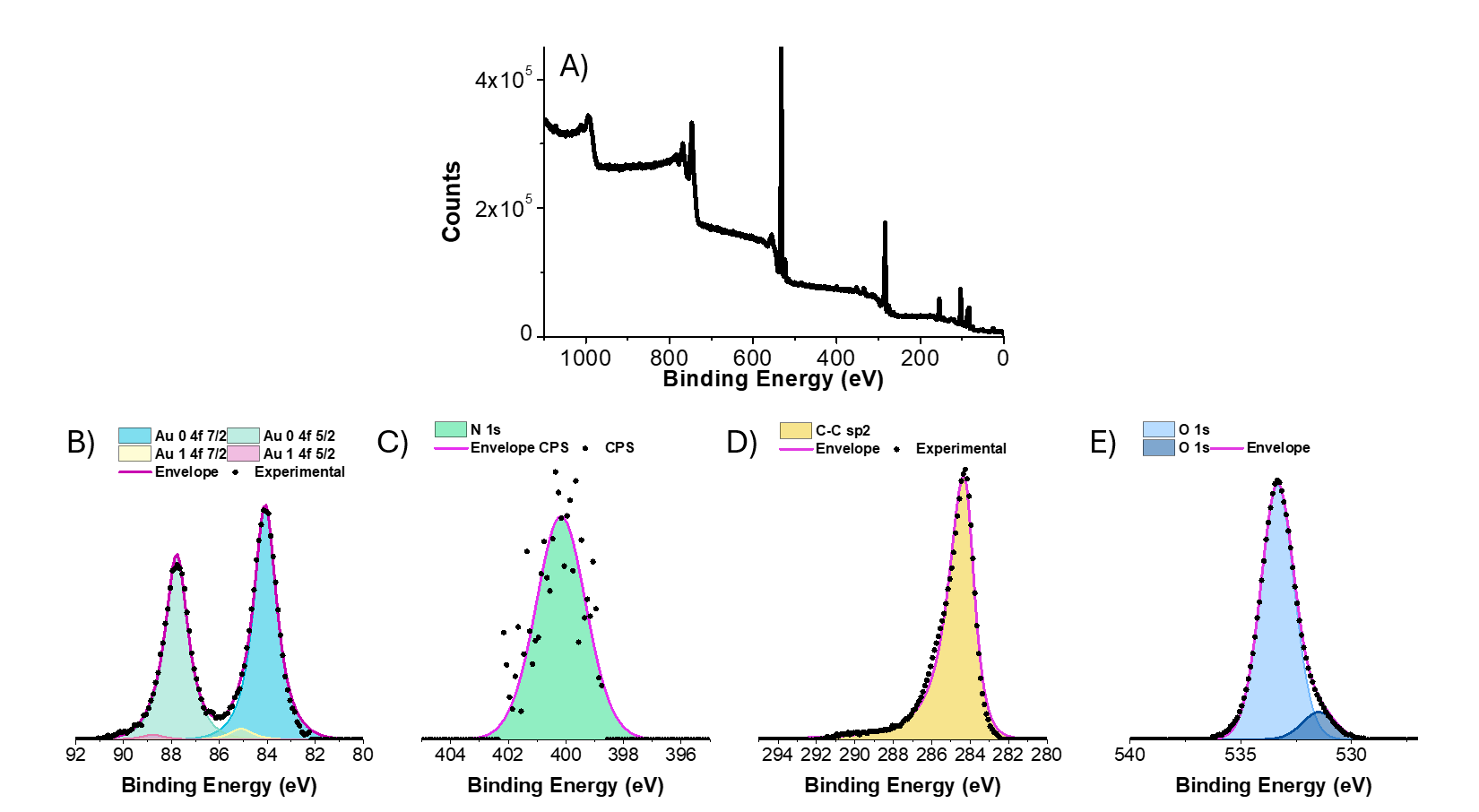

**Figure S2.** X-ray photoelectron spectroscopy (XPS) analysis of the Au-CNT composite. (A) Survey spectrum showing the characteristic core-level peaks of the constituent elements. (B) High-resolution XPS core-level spectra of Au 4f, showing the deconvoluted doublets corresponding to metallic gold Au^0^ (major component) and oxidized gold Au^1^ (minor component); (C) N 1s; (D) C 1s, dominated by the characteristic sp2-hybridized carbon (C=C) peak from the MWCNT scaffold; and (E) O 1s. Black dots represent experimental data points, and solid lines represent the fitted peak envelopes.

2. **Electrode surface characterization**

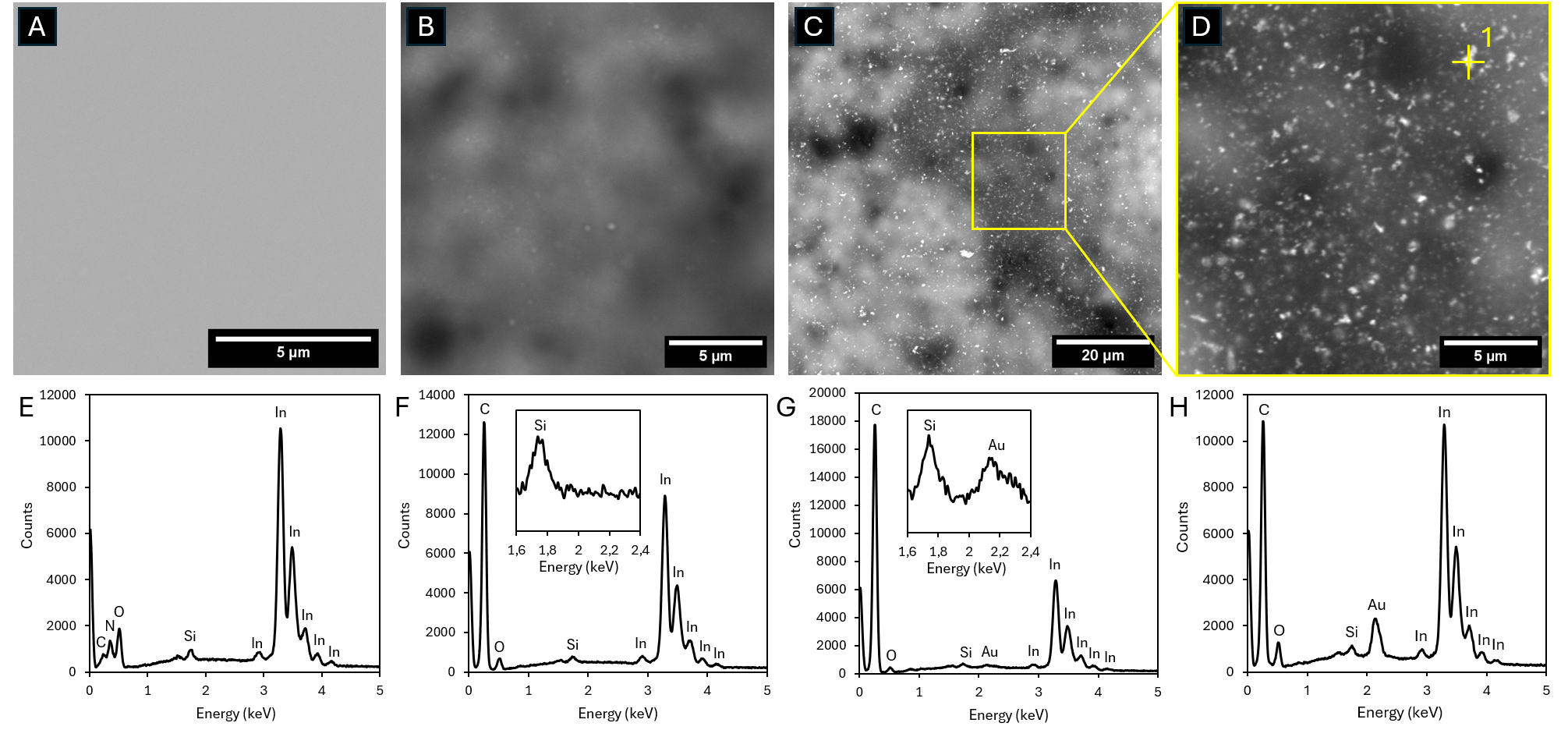

**Figure S3. Morphological and elemental characterization of electrode surface modifications. (A–D)** SEM micrographs and (**E–H)** corresponding EDX spectra recorded after each surface modification step: **(A, E)** Pristine ITO substrate; **(B, F)** ITO/CNT modified surface; **(C, G)** ITO/CNT**/**Au-CNT modified surface; **(D, H)** Magnified region of the ITO/CNT**/**Au-CNT modified surface showing targeted point-wise EDX analysis on a bright spot (Point 1 in D).

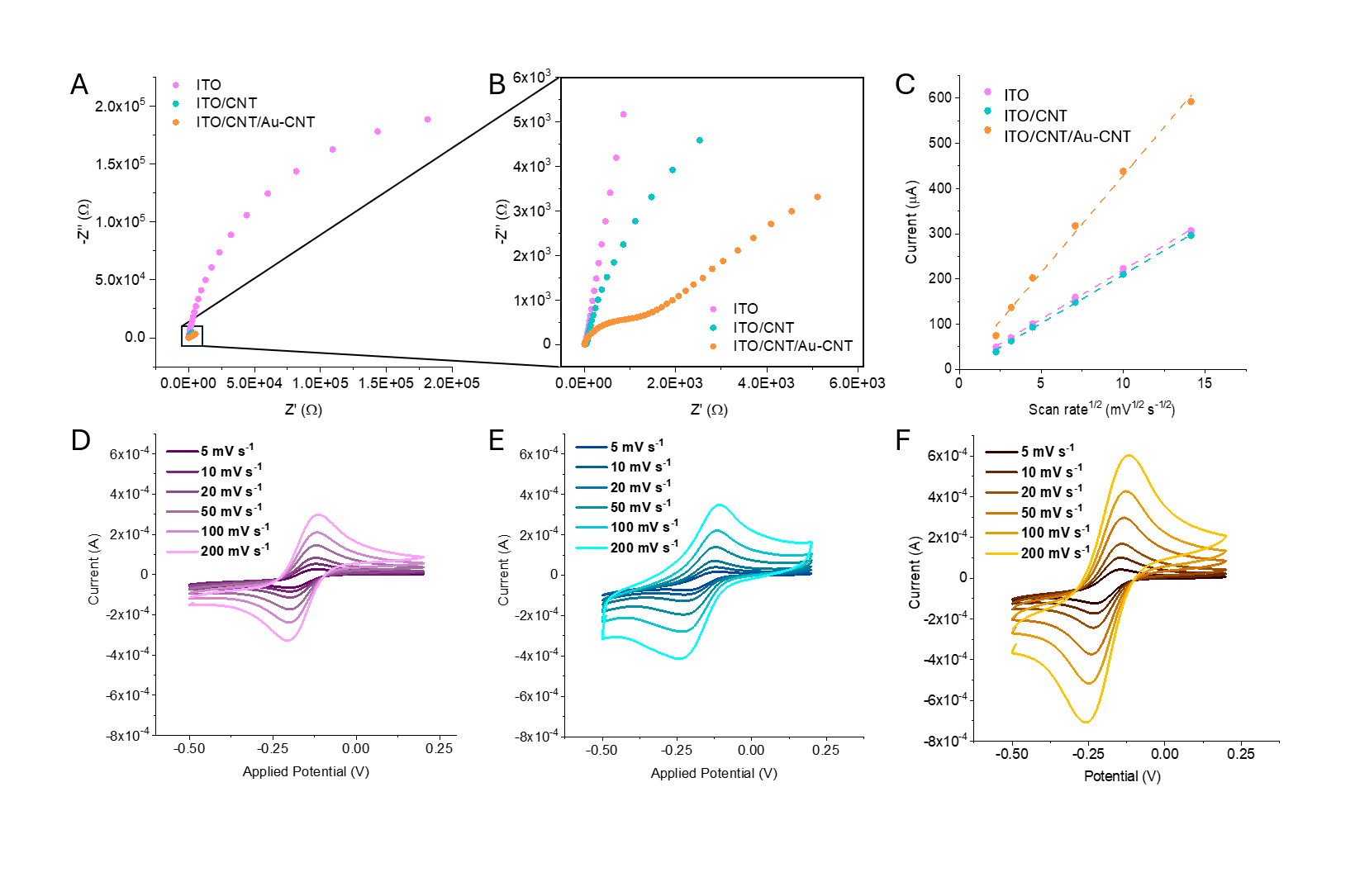

**Figure S4. Step-by-step electrochemical characterization of modified ITO electrodes.** (**A**) Nyquist plots obtained from Electrochemical Impedance Spectroscopy (EIS) in Krebs-Ringer buffer (pH 7.4) containing 100 𝜇M DA at E_1/2_ = 0.18 V vs. Ag/AgCl, with (**B**) an expanded view of the high-frequency region showing the low R_CT_ for modified electrodes. (**C**) Linear plots of anodic peak current versus the square root of scan rate (ν^1/2^) used for ECSA estimation. (**D-F**) CVs recorded in 1 μM [Ru(NH_3_)_6_]^3+^ in 0.1 M KCl at scan rates ranging from 5 to 200 mV s^-1^ for (**D**) pristine ITO, (**E**) ITO/CNTs, and (**F**) ITO/CNTs/Au-CNTs.

3. **Design of Experiment for the optimization of DPV parameters**

Differential Pulse Voltammetry (DPV) was chosen as an electroanalytical technique for the detection of DA, and a design of experiments (DoE) was used to optimize the system's performance. Two main factors were considered: modulation amplitude (MA), the intensity of each potential pulse, and modulation time (MT), the duration of each pulse. A factorial design was applied, choosing three equally spaced levels (0.01 V, 0.055 V, and 0.1 V for MA; 0.01 s, 0.055 s, and 0.1 s for MT) for each factor (**Table S1**).

**Table S1.** The two factors (MA and MT) and three levels chosen for the DoE.

|  | **MA (V)** | **MT (s)** |
| --- | --- | --- |
| **Low** | 0.01 | 0.01 |
| **Medium** | 0.055 | 0.055 |
| **High** | 0.1 | 0.1 |

Subsequently, all the possible combinations (n=9) were formulated, and each experiment was performed in triplicate with the same electrode (SPCE/Au-CNTs). The sequence of experiments was randomized for each series (**Table S2**).

**Table S2.** Sets of experiments for DoE.

| **SET 1** | | | **SET 2** | | | **SET 3** | | |
| --- | --- | --- | --- | --- | --- | --- | --- | --- |
| exp n. | MA (V) | MT (s) | exp n. | MA (V) | MT (s) | exp n. | MA (V) | MT (s) |
| 1 | 1 | -1 | 1 | -1 | 1 | 1 | 1 | -1 |
| 2 | 0 | -1 | 2 | 0 | 1 | 2 | 0 | -1 |
| 3 | 1 | 0 | 3 | 0 | 0 | 3 | 1 | 0 |
| 4 | 0 | 0 | 4 | -1 | 0 | 4 | 0 | 0 |
| 5 | 0 | 1 | 5 | 1 | 0 | 5 | 0 | 1 |
| 6 | 1 | 1 | 6 | 1 | 1 | 6 | 1 | 1 |
| 7 | -1 | -1 | 7 | 1 | -1 | 7 | -1 | -1 |
| 8 | -1 | 0 | 8 | 0 | -1 | 8 | -1 | 0 |
| 9 | -1 | 1 | 9 | -1 | -1 | 9 | -1 | 1 |

To evaluate the performance of the system, the following figure of merit was used:

**Equation 1**

$$FoM=\frac{i_{F}^{2}}{{\Delta E}_{1/2}i_{b}}$$

Where $i_{F}$ is the faradaic current, ${\Delta E}_{\frac{1}{2}}$ is the width of the peak at medium height, and $i_{b}$ is the baseline current sampled immediately before the start of the oxidation peak. After performing the experiments, average values for FoM were determined for each factor-level combination, as reported in **Figure S5.**

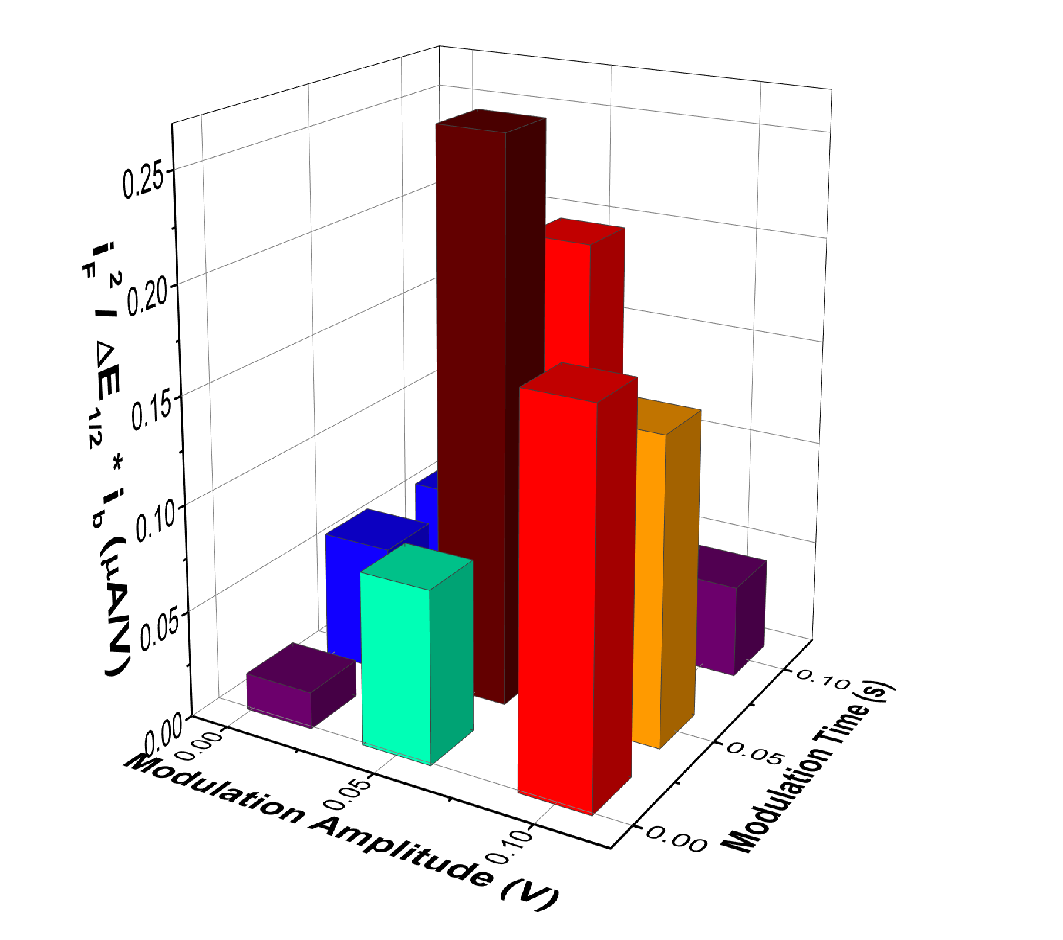

**Figure S5.** Average values obtained for the FoM from experimental data.

To obtain polynomial functions describing the dependence of FoM on MA and MT, a multiple regression analysis was performed. The following model was used to fit experimental data as a paraboloid:

$$FoM=b_{0}+b_{1} MA-b_{2} MT-b_{3} MA MT-b_{4} {MA}^{2}+b_{5}{MT}^{2}+b_{6}{MA}^{2}MT+b_{7}MA{MT}^{2}+b_{8}{MA}^{2}{MT}^{2}$$

The coefficient values are reported in **Table S3.**

**Table S3.** Coefficient values from regression.

| ***Regression coefficient*** | ***Corresponding term*** | ***Regression coefficient value*** |
| --- | --- | --- |
| *b_0_* | *Intercept* | 0.019 |
| *b_1_* | *MA (V)* | -2.31 |
| *b_2_* | *MT (s)* | -1.74 |
| *b_3_* | *MA MT* | 395.08 |
| *b_4_* | *MA^2^* | 39.17 |
| *b_5_* | *MT^2^* | 14.24 |
| *b_6_* | *MA^2^MT* | -3783.80 |
| *b_7_* | *MAMT^2^* | -2942.42 |
| *b_8_* | *MA^2^MT^2^* | 26714.51 |

The functions were normalized and combined in the FoM reported in Figure 1C.

4. **Study of the catalytic selectivity of Au-CNTs toward DA**
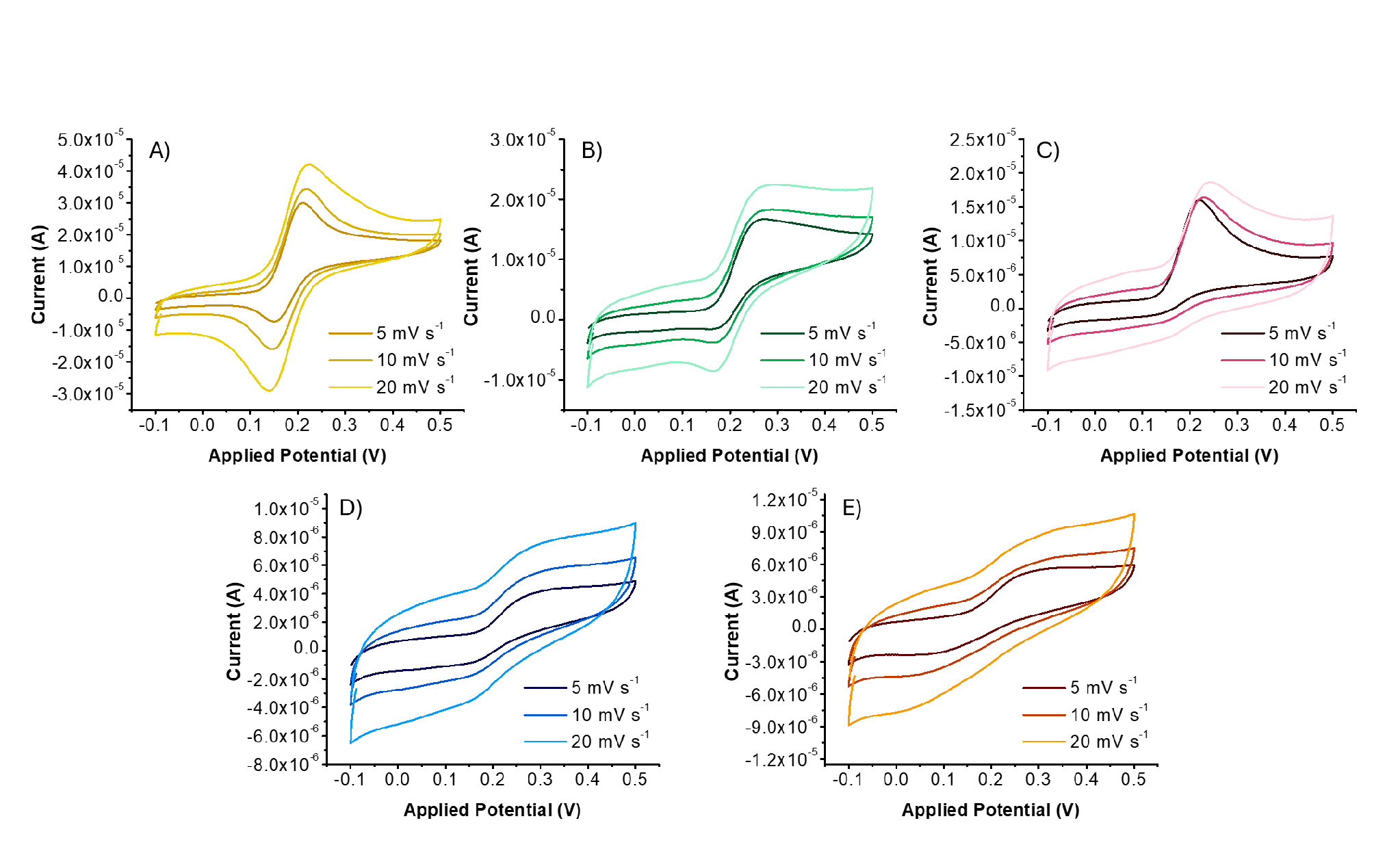

**Figure S6.** Voltammetric responses of 100 µM DA (A), NEP (B), EP (C), L-DOPA (D), and DOPAC (E) on ITO/CNT/Au-CNT electrodes in PBS 0.1 M and KCL 0.1 M at three different scan rates (5, 10, 20 mV s^-1^).
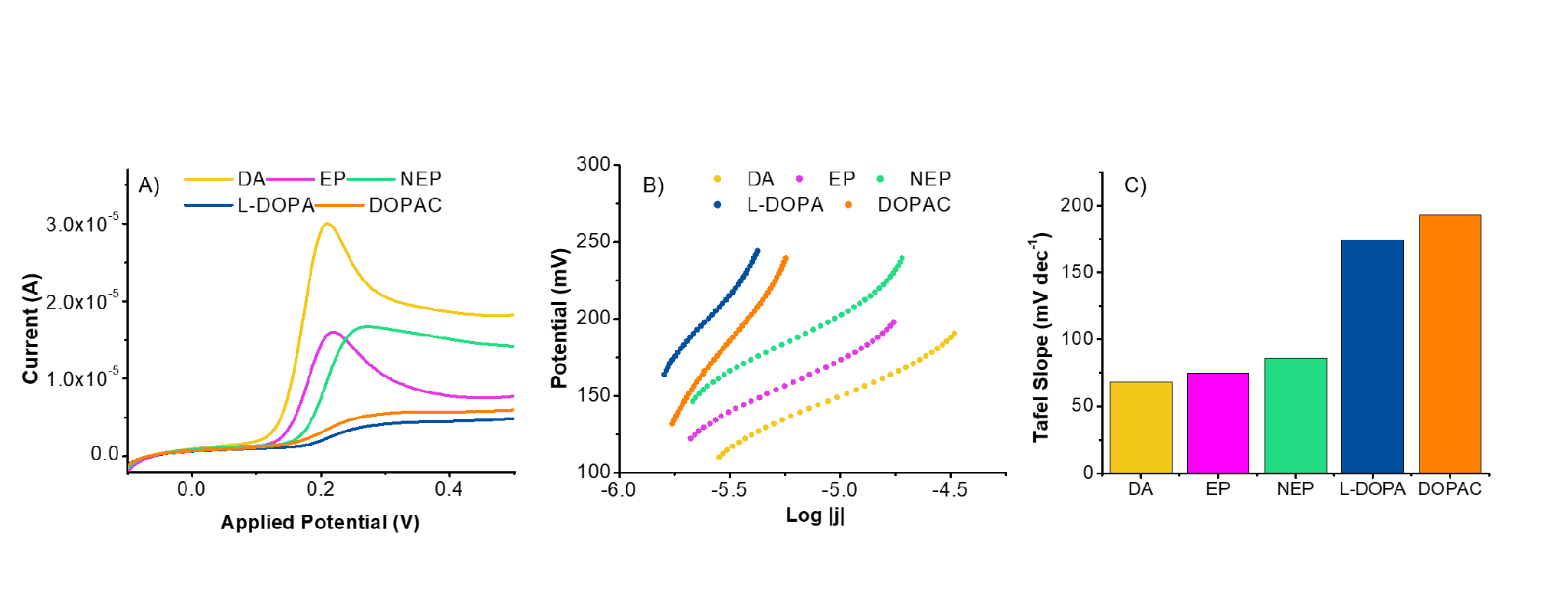

**Figure S7.** (A) Linear sweep voltammograms of 100 µM DA, EP, NEP, L-DOPA, and DOPAC on ITO/CNT/Au-CNT electrodes in 0.1 M PBS containing 0.1 M KCl at a scan rate of 5 mV s^-1^ (B) Tafel plots and (C) corresponding Tafel slopes for each analyte.

**Table S4.** ΔE and Tafel slopes recorded for the different neurotransmitters

| ***Neurotrasmitter*** | ***ΔE (mV)*** | ***Tafel Slope (mV/dec^-1^)*** |
| --- | --- | --- |
| *Dopamine (DA)* | *51* | *68.15* |
| *Norepinephrine (NEP)* | *80* | *85.85* |
| *Epinephrine (EP)* | *80* | *74.61* |
| *Levodopa (L-DOPA)* | *129* | *174.24* |
| *3,4-dihydroxyphenylacetic acid (DOPAC)* | *173* | *193.07* |

5. **Comparison of Au-CNT electrocatalytic properties with literature**

**Table S5.** Comparative table of electrochemical sensors for DA based on CNT/AuNP systems present in the state of the art.

| **Materials** | **AuNP synthesis** | **Method** | **E_ox_ peak** | **LOD** | **Linear range** | **Application** | **Ref.** |
| --- | --- | --- | --- | --- | --- | --- | --- |
| AuNBP/MWCNT / GCE (gold nanobipyramids on MWCNTs) | NaBH_4_/Citrate reduction (capped) | DPV | 0.24 V (CV) | 600 nM | 0.2 – 157 μM | Buffer / spiked human fluids | [1] |
| Au@NAC-MWCNT / GCE (NAC-stabilized Au clusters) | Hydrothermal (NAC-capped) | DPV/ CV | ~0.17 V (CV) ~0.15 V (DPV) | 30 nM | 0.1 – 250 µM | Spiked serum/urine | [2] |
| GNPs/MWCNTs / GCE (gold nanoparticle on MWCNTs) | Modified Brust-Schiffin, NaBH_4_ reduction (Capped) | SWV | ~0.30 V (CV) | 71 nM | 0.48 - 5.7 μM | Buffer, synthetic cerebrospinal fluid | [3] |
| N-MWCNT/AuNPs / film on Si wafer (AuNP on Nitrogen-doped MWCNTs) | Photochemical segmented flow, (PVP-capped) | SWV | 0.237 V | 1.60 μM | 10 - 350 μM | Diluted pig serum | [4] |
| AuNPs/DWCNT/ GCE (gold nanoparticles on DWCNTs) | Electrodeposition | CV | 0.252 V (vs SCE) | 1.3 μM | 13 - 163 μM | Buffer | [5] |
| AuNPs/CNT-PCA / GCE (pyrene carboxylic acid linker) | Citrate reduction (capped) | DPV | ~0.16 V (CV) | 1 nM | 100 nM – 150 µM | Spiked human blood serum | [6] |
| AuNPs-CNT/ITO | MVS | LSV/ DPV | 0.20 V (CV) | 412 nM | 0.8-7 µM | Krebs-Ringer buffer | This work |

6. **Reproducibility and repeatability**

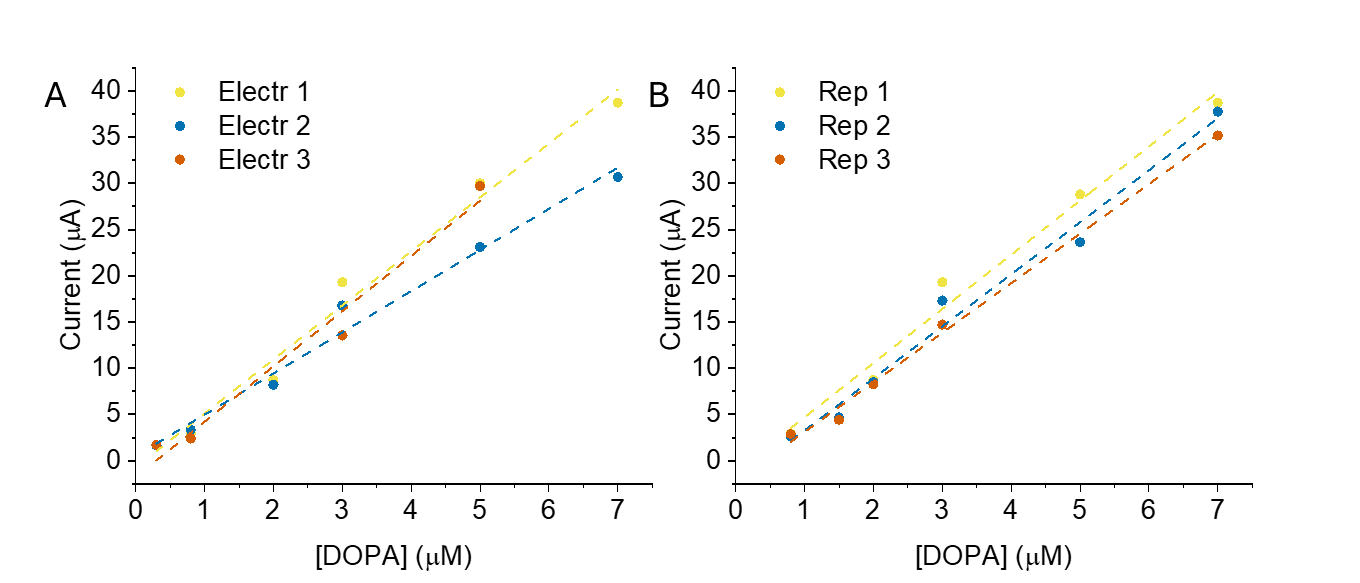

**Figure S8. Assessment of sensor repeatability and reproducibility.** (**A**) Reproducibility evaluated across three independently fabricated ITO/CNT/Au–CNT-modified electrodes under identical experimental conditions. Linear dashed lines represent the corresponding calibration fits. (**B**) Repeatability evaluated by performing three consecutive calibration curves on a single ITO/CNT/Au–CNT-modified electrode across the linear range.

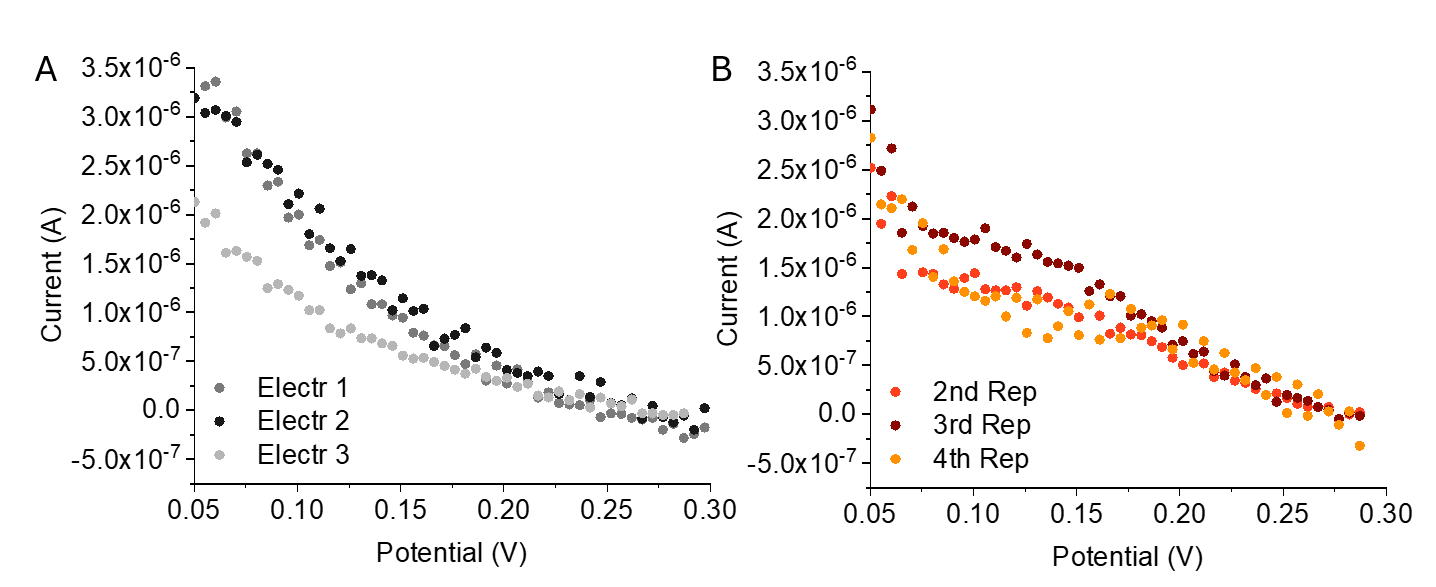

**Figure S9.** Baseline measurements in buffer recorded on (**A**) three fresh electrodes and (**B**) three electrodes reused after a calibration run and subjected to buffer washing and electrochemical cleaning (5 CV cycles between 0.0 V and 0.5 V at a scan rate of 50 mV s^-1^), demonstrating minimal residual dopamine/polydopamine signal.

7. **Long-term stability of the electrodes**

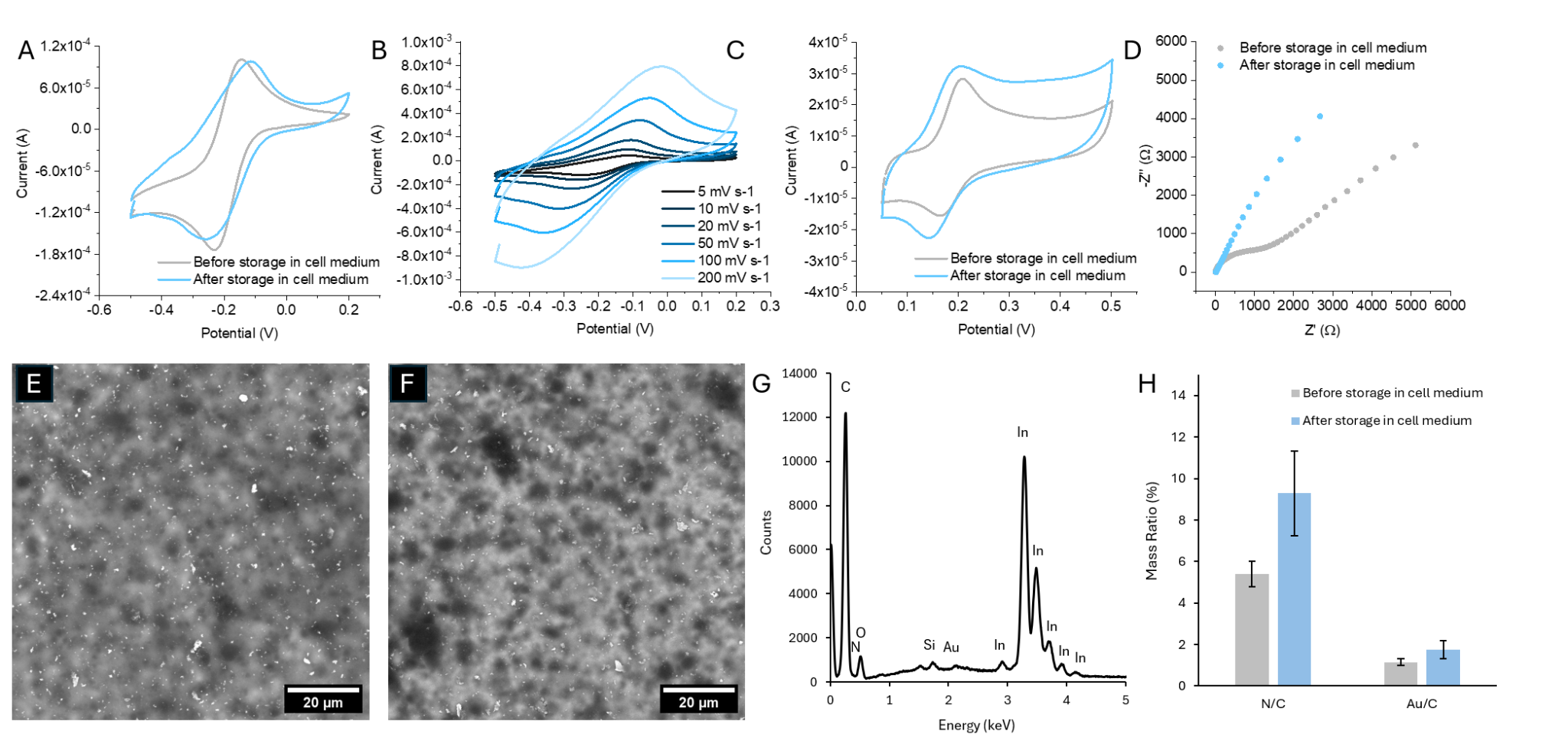

**Figure S10. Surface and electrochemical characterization of the modified electrode before and after 21 days of incubation in cell culture medium.** **(A)** CV curves recorded in 1 μM [Ru(NH_3_)_6_]^3+^ before (grey) and after (blue) incubation; **(B)** CVs of the post-incubated electrode in [Ru(NH_3_)_6_]^3+^ recorded at various scan rates (5-200 mV s^-1^) highlighting the increase in ohmic drop with the scan rate; Cyclic voltammograms **(C)** CV recorded in Krebs medium with 100 μM dopamine before (grey) and after (blue) 21 days of storage in cell medium; **(D)** Nyquist plots obtain in 100 μM dopamine before (grey) and after (blue) storage; **(E, F)** Topographical SEM images of the electrode surface **(E)** before and **(F)** after 21 days of incubation in cell medium; **(G)** EDX spectra of the electrode surface after 21 days of storage in cell medium; **(H)** EDX elemental mass ratios (N/C and Au/C) before (grey) and after (blue) incubation.

8. **iPSC induction to DAergic neurons**

**Table S6.** DAergic neuron conversion media recipe.

| **NC Medium [Neuron Conversion Media]** | | |
| --- | --- | --- |
| **DMEM:F12/Neurobasal-Brainphys (1:1)** | | |
| **Supplements** | **Cf** | **Source** |
| **N-2 Supplement (100X)** | 1X | Fisher Scientific |
| **B-27™ Supplement (50X), minus Vitamin A** | 1X | Fisher Scientific |
| **Doxycycline Hyclate (WC2031)** | 2 µg mL^-1^ | Selleckchem |
| **Laminin from Engelbreth-Holm-Swarm murine sarcoma** | 1 µg mL^-1^ | Sigma Aldrich |
| **Gibco™ DMEM/F-12, GlutaMAX™ supplement** | 100X | Thermo Fisher |
| **LDN-193189 hydrochloride** | 200 nM | Sigma Aldrich |
| **Recombinant Mouse Sonic Hedgehog/Shh (C25II) N-Terminus** | 100 ng mL^-1^ | RDsystems |
| **Purmorphamine** | 2 µM | Sigma Aldrich |
| **C-6556 CHIR99021, Free Base, >99%** | 2.5 µM | Sigma Aldrich |
| **SB-431542** | 10 µM | MedChem |

**Table S7.** DAergic neuron maturation media recipe.

| **NM Medium [Neuron Maturation Media]** | | |
| --- | --- | --- |
| **DMEM:F12/Neurobasal-Brainphys (1:1)** | | |
| **Supplements** | **Cf** | **Source** |
| **N-2 Suplement (100X)** | 1X | Fisher Scientific |
| **B-27™ Plus Suplement (50X)** | 1X | Fisher Scientific |
| **Doxycycline Hyclate (WC2031)** | 2 µg mL^-1^ | Selleckchem |
| **Laminin from Engelbreth-Holm-Swarm murine sarcoma** | 1 µg mL^-1^ | Sigma Aldrich |
| **Dibutryryl cAMP** | 0.10 mM | StemCell |
| **Non essential amino acids MEM (100X)** | 100X | Thermo Fisher |
| **Insulin-Transferrin-Selenium (ITS-G)** | 1X | Thermo Fisherch |
| **GlutaMAX™ supplement** | 100X | Thermo Fisher |
| **Recombinant Human TGF-β3** | 1 ng mL^-1^ | Sigma Aldrich |
| **DAPT** | 10 µM | Selleckchem |
| **GDNF, Human Recombinant  Animal Free** | 20 ng mL^-1^ | Sigma Aldrich |
| **Brain Derived Neurotrophic Factor (BDNF)** | 20 ng mL^-1^ | MedChem |

9. **iPSC induction to GLUTergic neurons**

**Table S8**. GLUTergic neuron conversion media recipe.

| **Neuron Conversion Media** | | |
| --- | --- | --- |
| **DMEM:F12/Neurobasal-Brainphys (1:1)** | | |
| **Supplements** | **Cf** | **Source** |
| **N-2 Suplement (100X)** | 1X | Fisher Scientific |
| **B-27™Plus Suplement (50X)** | 1X | Fisher Scientific |
| **Doxycycline Hyclate (WC2031)** | 2 µg mL^-1^ | Selleckchem |
| **Laminin from Engelbreth-Holm-Swarm murine sarcoma** | 1µg mL^-1^ | Sigma Aldrich |
| **Dibutryryl cAMP** | 100 µg mL^-1^ | stemCell |
| **Human Recombinant Noggin** | 150 ng mL^-1^ | stemCell |
| **LDN-193189 hydrochloride** | 0.5 µM | Sigma Aldrich |
| **A83-01** | 0.5 µM | Selleckchem |
| **Forskolin** | 5 µM | MedChemExpress |
| **C-6556 CHIR99021, Free Base, >99%** | 3 µM | Sigma Aldrich |
| **SB-431542** | 10 µM | MedChem |

**Table S9.** GLUTergic neuron maturation media recipe.

| **Neuron Maturation Media** | | |
| --- | --- | --- |
| **DMEM:F12/Neurobasal-Brainphys (1:1)** | | |
| **Supplements** | **Cf** | **Source** |
| **N-2 Supplement (100X)** | 1X | Fisher Scientific |
| **B-27™ Plus Supplement (50X)** | 1X | Fisher Scientific |
| **Doxycycline Hyclate (WC2031)** | 2 µg mL^-1^ | Selleckchem |
| **Laminin from Engelbreth-Holm-Swarm murine sarcoma** | 1 µg mL^-1^ | Sigma Aldrich |
| **Dibutryryl cAMP** | 100 µg mL^-1^ | StemCell |
| **GDNF, Human Recombinant  Animal Free** | 20 ng mL^-1^ | Sigma Aldrich |
| **Brain Derived Neurotrophic Factor (BDNF)** | 20 ng mL^-1^ | MedChem |

**Table S10.** qPCR primers: forward and reverse sequences.

| **Target** | **Forward** | **Reverse** |
| --- | --- | --- |
| **Ascl1** | 5’-GAATGGACTTTGGAAGCAG | 5’-TTTCTTTTCCTTTTCTCCCC |
| **Ngn2** | 5’-AAGATGTTCGTCAAATCCG | 5’-CTCCTCCTCCTCTTCTTC |
| **TUBB3** | 5’-TCATCTTTGCTCAGACTGG | 5’-GTTTTCACACTCCTTCC |
| **TH** | 5'-CAAAATCCACCATCTAGAGAC | 5’-TGACACTTTTCTTGGGAAC |
| **GAD1** | 5'-CAATACCAACATCCCATCAG | 5’-ATTTAGGAGGAAGACCCTTG |
| **vGLUT1** | 5’-GCAAGTACATCGAGGACGCCAT | 5′-TCAGTAGTCCCGGACAGGGG |
| **CHAT** | 5’-TCAGTTCTTTGTCTTGGATG | 5’-TGGAAGCCATTTTGACTATC |
| **GFAP** | 5'-CAAGATGAAACCAACCTGAC | 5’-CTTCCTCCTCATGGATCTTC |
| **GAPDH** | 5 ́-ACAGTTGCCATGTAGACC | 5 ́-TTGAGCACGGGTACTTTA |
| **Actin** | 5’-CTCTATCATGAAGTGCGATG | 5’-CCCCAATCCAAACAGAATAC |

10. **Viability assays**

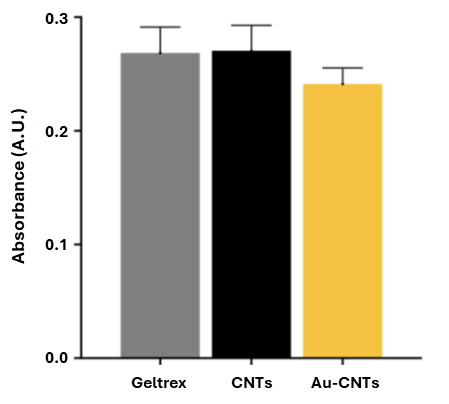

**Figure S11.** Viability of iPSCs measured by LDH assay in vitro in Geltrex, CNTs, and Au-CNTs. Bar chart showing the absorbance at 490nm. Absorbance values were obtained after 5 days of iPSC cell culture with an initial seeding density of 100,000 cells per well. The statistical significance was assessed using t-tests. Error bars represent the SD of four replicates (mean ± SD, n = 4)

11. **Measurement in cellular cultures**

**Table S11.** Comparative table of basal endogenous levels typically observed in PC12 cell cultures

| **Cell Type** | **LOD** | **Linear Range** | **Electrochemical Technique(s)** | **Dopamine Concentration** | **Ref.** |
| --- | --- | --- | --- | --- | --- |
| SH-SY5Y cells, matured into DAergic neurons | 33.4 nM | 0.5-100 μM | DPV | - | ^[7]^ |
| PC12 | 12  nM | 50 nM to 288 μM | CA | - | ^[8]^ |
| PC12 | 2  nM | 5 nM to 1 μM | DPV-CA | 48.5 nM | ^[9]^ |
| PC12 | 1.279 nM | 0-250  nM | CV-CA | 43.1 ± 21.6 nM | ^[10]^ |
| Differentiated human neural stem cells | - | - | CA | - | ^[11]^ |
| SH-SY5Y | 60  nM | 1-200  μM | DPV | - | ^[12]^ |
| PC12 | 0.87 nM | 0.005 to 100.0 μM | DPV | No signal detected in the absence of KCl | ^[13]^ |
| PC12 | 134  nM | 0.1-50  μM | DPV | - | ^[14]^ |

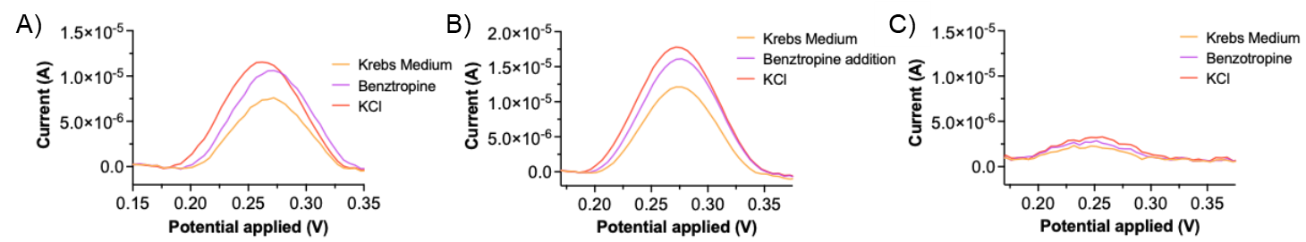

**Figure S12.** DPV was recorded in the DAergic neuron before and after the addition of Benzotropine and KCl.

12. **Cell-free control experiments**

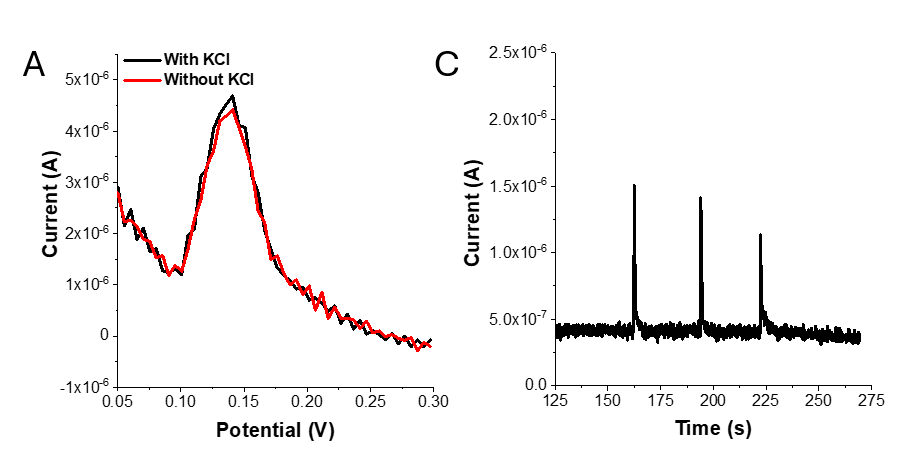

**Figure S13. Cell-free electrochemical controls assess the influence of KCl addition. (A)** Baseline-subtracted DPV voltammograms of 1.5 μM dopamine in Krebs-Ringer buffer recorded before (red line) and after (black line) the addition of KCl (80 μM final concentration increment). **(B)** Chronoamperometric current-time trace (0.27 V vs. Ag/AgCl) showing three consecutive additions of 10 μL of a 1.6 mM KCl stock solution to 200μL of a 1.5 μM dopamine solution in Krebs-Ringer buffer. The sharp, transient current spikes correspond to fluid turbulence induced by pipetting, with the current returning to the baseline.
